# Antigen avidity drives the durability of the vaccine immune response

**DOI:** 10.1101/2025.04.18.649567

**Authors:** Nicole G. Weidner, Mauricio V. Padilla, Layne Pruitt, Christopher A. Cottrell, Kristyn Gonzales, Oleksandr Kalyuzhniy, Michael Kubitz, Nushin Alavi, Nicole Phelps, Meredith Weglarz, William R. Schief, Robert K. Abbott

**Affiliations:** Department of Pathology, University of Texas Medical Branch, Galveston, TX, USA; Department of Immunology and Microbiology, The Scripps Research Institute, La Jolla, CA; IAVI Neutralizing Antibody Center, The Scripps Research Institute, La Jolla, CA 92037, USA; Center for HIV/AIDS Vaccine Development, The Scripps Research Institute, La Jolla, CA 92037, USA; Moderna Inc., Cambridge, MA 02139, USA

## Abstract

One challenge in vaccine development is designing immunogens that elicit durable immunity. We hypothesized that antigen avidity regulates the magnitude, diversity, and durability of the vaccine immune response. We tested this in multiple preclinical HIV vaccine models using a neoteric mosaic nanoparticle platform. This allowed us to precisely modulate antigen avidity by varying multivalency and affinity independently, whilst keeping other variables constant. Antigen avidity drove seeding, interclonal competition, and immunodominance within germinal centers. High-valency immunogens promoted durable germinal center, memory B cell, serum antibody, and long-lived plasma cell responses. Restricting interclonal competition rescued B cell responses to low-valency immunogens. B cell receptor sequencing revealed that antigen valency had minimal impact on individual somatic hypermutations but promoted clonal diversity. Affinity worked in concert with valency in driving productive B cell responses, with multivalency having dominant influences. The results underscore the importance of antigen avidity in driving durable and diverse vaccine responses.

## Introduction

Vaccines are arguably the greatest medical invention of the 20^th^ century, saving an estimated 154 million lives over the last 50 years.^1^ Moreover, vaccines are incredibly safe and effective, offering one of the highest risk to reward ratios of any medical intervention. However, vaccines have been a victim of their own success. Our foundational mechanistic knowledge surrounding what critical components of a vaccine drive durable immune responses remains incomplete. This is in part due to a lack of truly predictive mechanistic preclinical models that recapitulate the challenges that exist in humans. One of these challenges includes developing models that match the human precursor frequency of targeted B cells that can develop protective neutralizing responses. Another is the utilization of complex immunogens that elicit B cell responses that mimic the high levels of interclonal competition that exists in response to real world vaccines.

Antigen avidity is the generic phrase coined for the overall strength of the antigen-B cell receptor (BCR) interaction. Antigen avidity is determined by the product of two principal components – epitope affinity and epitope valency. While affinity is purely the monomeric binding strength of individual epitope-Fab interactions, valency is the overall level of repetitiveness of the epitope. Multiple preclinical studies have suggested that antigen valency is likely an important driver of successful B cell responses post vaccination^2–10^, but open questions remain. Antigen affinity has also been shown to be independently important in selectively priming desired B cell responses following vaccination both preclinically^8^ and clinically^11^. Increased antigen valency is generally considered to promote a more robust serum antibody response post vaccination in humans.^12,13^ This general observation has sparked development and deployment of highly valent nanoparticle-based and virus-like particle (VLP) based vaccine platforms. This includes clinical and preclinical development for multiple targets including hepatitis B,^14^ human papilloma virus,^15^ influenza,^16–19^ respiratory syncytial virus,^20,21^ SARS-CoV-2,^22–27^ and HIV.^9,11,28^ Longitudinal studies in humans revealed that the half-life of serum antibody responses to low-valency vaccines such as tetanus and diphtheria was substantially less than high-valency vaccines (e.g.; measles, mumps, rubella, and vaccinia).^12^ However, these vaccines differ not only in the valency of the antigens, but also in adjuvanticity, replicative capacity, immunogen size, and T cell help epitope qualities. This makes direct conclusions about antigen valency driving vaccine durability difficult. Are some of these vaccines more durable due strictly to increased antigen valency, or due to other variables, or a combination of variables? More definitive mechanistic studies are needed to tease apart the critical individual components of successful vaccines that drive durability. There is currently no accepted guidance on minimum epitope valencies that should be used for human subunit vaccines. Human clinical trials are still initiated with low-valency vaccine immunogens. This includes candidate vaccines for West Nile virus,^29^ dengue^30^ and HIV.^31^

Recently, antigen valency has been utilized in next generation mosaic vaccine designs to recruit broadly protective B cells for multiple viral targets including SARS-CoV-2^24–26^ and influenza.^17^ This has been done by utilizing antigen valency as a rheostat to preferentially recruit cross-reactive B cell responses. In this approach, mosaic nanoparticles contain mixed protective epitopes from different viral strains on the same particle. This tested the prediction that only B cells with broad strain reactivity will have a competitive advantage presumably due to increased BCR crosslinking relative to non-cross reactive B cells. While this was observed in the serological response, the cellular mechanisms by which mosaic nanoparticles actually work remains unknown.

Germinal centers (GCs) are specialized structures within lymph nodes that are the engines of vaccine immune responses.^32–34^ Following vaccination, antigen activated B cells proliferate and undergo T cell-mediated competition^35,36^ to enter GCs within days. B cells proliferate rapidly within GCs and undergo somatic hypermutation (SHM) of their antibody-encoding genes to develop improved antibody responses. High affinity B cell clones are preferentially selected by T follicular helper cells (Tfh)^37–39^ within light zones of GCs to return to dark zones to proliferate.^40–42^ Taken together, the process of mutation and selection has been coined the “cyclic re-entry”^43^ model of clonal selection. This process gives mechanistic explanation to the affinity maturation process. Affinity maturation is the phrase coined to describe the stepwise orders of magnitude increases of serum antibody affinity in the months following vaccination.^33,44^ At the cellular level, successful B cells exit GCs as one of two enduring fates. The first is as memory B cells (MBCs) that can last a lifetime^45^ and rapidly respond to reinfection.^46^ The second are long-lived plasma cells (LLPCs) that reside in the bone marrow (BM). LLPCs secrete affinity matured antibodies into the serum for upwards of a lifetime following vaccination.^12,13,47–49^

The discovery of potent broadly neutralizing antibodies (bnAbs) to HIV^50^ provided renewed hope an HIV vaccine is possible. However, bnAbs develop only after years of infection^51–53^ and often display high levels of SHM through complex affinity maturation pathways.^54–56^ Interclonal competition post-vaccination affects the ability of bnAb-precursor B cells to enter and compete within GC reactions and exit as durable MBCs and LLPCs.^57,58^ To activate desired bnAb-precursor B cells, modified HIV envelope (Env) immunogens targeting the germline bnAb-precursor B cells have been developed,^9–11,59,60^ as bnAb precursors do not readily bind mature HIV Env.^61^ VRC01-class bnAbs neutralize HIV by binding the functionally essential CD4 binding site (CD4bs) epitope on Env, and are particularly attractive for vaccine design for multiple reasons. VRC01-class bnAbs are highly potent,^50^ safe,^62^ and have an HCDR2 dominant binding modality that is amenable to germline targeting.^63^ Moreover, a large percentage of humans are estimated to have VRC01-class bnAb precursor B cells^64^ in which early clinical trials have shown can be primed by germline targeting immunogens.^11^ One set of VRC01-class germline-targeting priming immunogens have been coined “<u>e</u>ngineered <u>o</u>uter-<u>d</u>omain <u>g</u>ermline targeting” (eOD-GT) immunogens.^9^

Several preclinical mouse B cell transfer systems using knock-in B cells expressing human heavy and light chain immunoglobulins have been developed to evaluate HIV immunogens under physiological conditions.^8,65–68^ These systems utilize an allotype marked B cell transfer approach coupled with precisely engineered immunogens that match human B cell precursor frequencies and antigen affinities.^64^ The VRC01^gHL^ system makes use of B cells containing germline reverted heavy and light chain sequences of the human broadly neutralizing antibody, VRC01.^8^ Next generation B cell transfer systems were developed utilizing authentic precursor B cell sequences that were recovered from healthy human donors^64,65^ and coined Human Germline (HuGL)^65^ mice.

Initial studies with the VRC01^gHL^ B cell transfer system revealed that some level of valency was required to prime rare VRC01^gHL^ B cells during vaccination from physiologically relevant precursor frequencies and affinities.^8^ This was evident as monomeric versions of immunogens were only able to prime VRC01^gHL^ B cells to enter GCs at supraphysiological pico-molar affinities, while 60-mers were able to within the human physiological range.^8^ However, a natural limitation of this interpretation was that 60-mer based nanoparticles are of both larger size (∼2400kDa vs ∼20kDa) and contain more T cell help epitopes. This is in part because the monomeric immunogens lack the lumazine synthase (LS) component that drives nanoparticle formation. Follow-on work utilizing this system found that priming of VRC01^gHL^ B cells with high-valency nanoparticle vaccine immunogens, compared to smaller, low-valency immunogens, facilitated stronger early B:T cell interactions, early GC and plasma cell (PC) differentiation, and GC competition.^6^ However, induction of MBCs, LLPCs, or vaccine durability was not assessed in that study. Moreover, one limitation of that study was that in order to address the challenge that different sized immunogens also contain differential quantity and quality of T cell help epitopes, the SMARTA peptide gp_61-80_ was added to each immunogen and SMARTA-1 CD4 T cells were co-transferred with each experimental condition (1mer, 4mer, 8mer, 60mer). This was done to attempt to normalize T cell help across different sized particles with different levels of MHCII I-A^b^ restricted helper epitopes.^6^

In the current study, we circumvented previous challenges of evaluating the role of antigen valency in driving durable vaccine immune responses to complex real-world immunogens. We achieved this by developing mixed mosaic nanoparticle immunogens, coupled with evaluation in multiple established preclinical mouse models for HIV vaccine development. This was done with both the preclinical eOD-GT5^8,9^ 60mer and clinically validated eOD-GT8^11,64^ 60mer immunogen platforms coupled with the VRC01^gHL^, HuGL17, and HuGL18 B cell transfer systems. These models utilized defined B cell precursor frequencies and affinities and were predictive of the phase 1 clinical trial of the HIV immunogen eOD-GT8 60mer.^11^ These highly controlled systems allowed us to test the hypothesis that antigen avidity, and particularly multivalency, was critical in driving both successful and durable vaccine immune responses.

## Results

### Antigen valency drives seeding, clonal bursting, and durability of germinal centers

To study the impact of B cell epitope valency in driving successful GC responses and durable immunological memory, we developed a LS-based mosaic nanoparticle immunogen system using our eOD-GT immunogens **(Fig. 1A-B)**. This system features two types of subunits: “wild type” (WT) and “knock out” (KO), which differ exclusively at the VRC01-class CD4 binding site (CD4bs) epitope due to only four point-mutations (N280R, S365L, F371R, D368R) that essentially ablates VRC01-class precursor binding. This allows us to precisely vary B cell epitope avidity by controlling the B cell epitope valency. Importantly, the WT and KO subunits retain other conserved regions, allowing us to vary B cell epitope valency while maintaining constant T cell helper epitopes and preserving immunogen size. The WT and KO subunits are each fused to the LS core subunits independently. Upon co-transfection they self-assemble into nanoparticles containing 60 total copies of a protein epitope. The 60 copies consist of a mixture of WT and KO subunits, producing a mosaic within each nanoparticle. This provides a robust system to investigate the role of epitope valency in vaccine immune responses.

**Figure 1:**
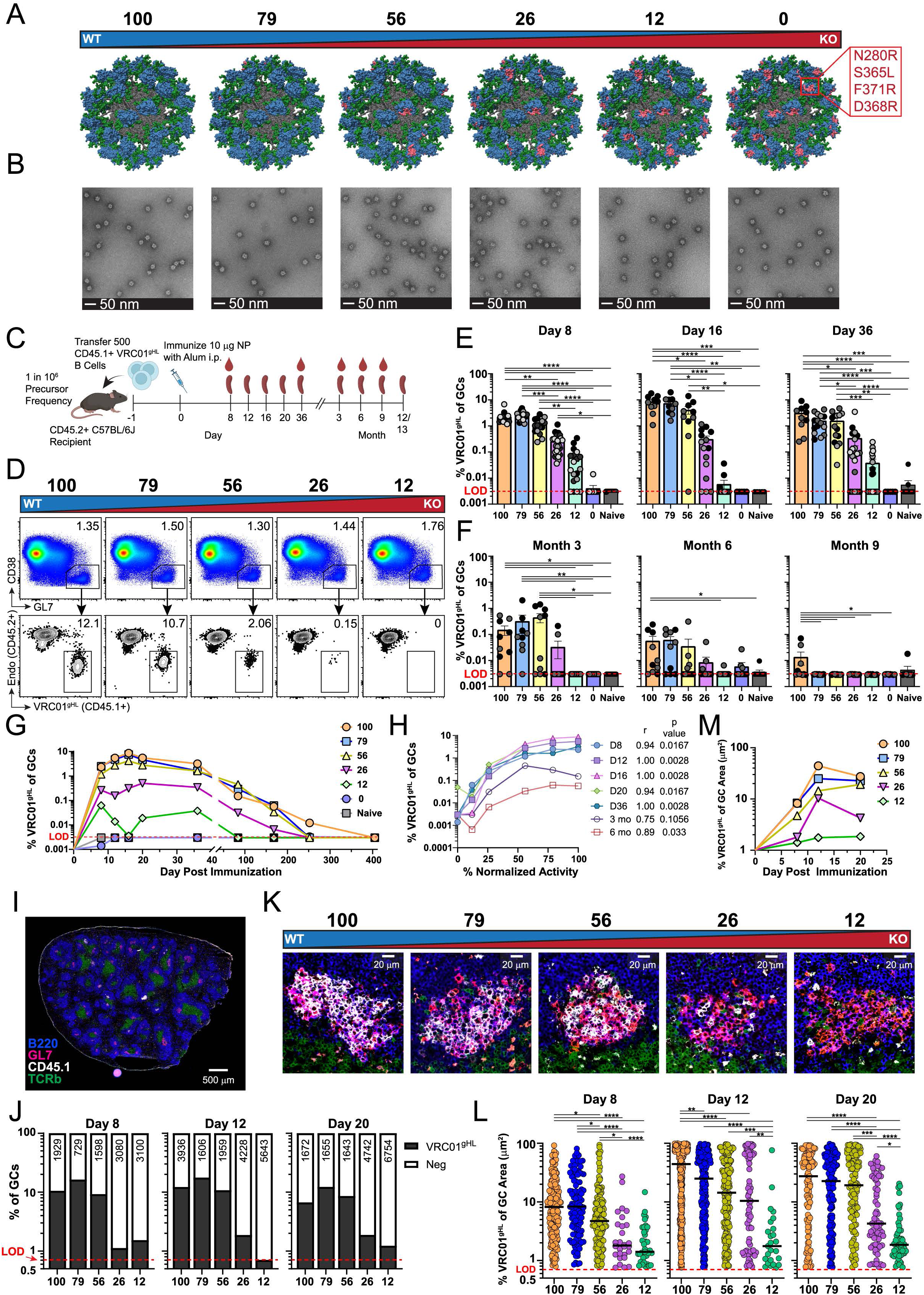
Antigen valency drives seeding, clonal bursting, and durability of germinal centers. **A)** Mosaic nanoparticles were developed with varying ratios of GT5 WT to GT5 KO. **B)** Negative stain EM images of mosaic nanoparticles listed in A. **C)** Schematic of the VRC01^gHL^ B cell transfer system. **D)** Representative flow cytometry from day 16 for quantifying VRC01^gHL^ B cells (CD45.1+) against endogenous cells (CD45.2+) within total GCs (CD38-GL7+) from mice immunized with mosaic nanoparticles. Pre-gated on SSL/B220+CD4-. **E-F)** Frequency of VRC01^gHL^ GC B cells gated as in D for **(E)** days 8, 16, and 36, and **(F)** months 3, 6, and 9. **G)** Frequency of VRC01^gHL^ B cells within GCs over time as shown in E and F. **H)** Spearman coefficient analysis for correlation between mosaic nanoparticle normalized activity and mean VRC01^gHL^ GC occupancy at the timepoints indicated. **I)** Representative spleen GC histology from day 12. **J)** Quantification of the percent of total GCs analyzed (number in graph) that were either positive or negative for VRC01^gHL^ B cells (CD45.1+). n = 5 mice/group. **K)** Representative splenic GCs on day 12 from mice immunized with different mosaic nanoparticles. **L)** Quantification of the percent area of VRC01^gHL^ in each GC positive for VRC01^gHL^ cells as shown in K. n= 5 mice/group. Bar indicates median. **M)** Quantification of average GC area occupied by VRC01^gHL^ B cells over time. n= 5 mice/group. E = independent experiment. **E-G)** E = 1-5. **H-L)** E = 1-2. **E-F, L)** Kruskal-Wallis with Dunn’s correction. *p<0.05, **p<0.01, ***p<0.001, ****p<0.0001. **E-F)** Mean ± SEM. Shades of grey dots indicate independent experiments. **E-G, J-L)** LOD = limit of detection. See also Figure S1 and Table S1.

Our first studies of these mosaic-based nanoparticles were done with the immunogen eOD-GT5 60mer in the VRC01^gHL^ model system **(Fig. 1C)**. In the VRC01^gHL^ model system,^8^ the earlier generation eOD-GT5 60mer is used to mimic the affinity range of true human naïve VRC01-class precursors^64^ for the clinically validated immunogen eOD-GT8 60mer.^11^ The affinity of eOD-GT5 for VRC01^gHL^ is in the higher range of known physiologically relevant affinities of human VRC01-class precursors (∼0.2 to 0.5µM *K*_D_).^6,8,9^ We generated four variants of eOD-GT5 60mer by co-transfecting HEK293 cells with varying ratios of WT and KO GT5 constructs to form particles with mixed subunits **(Table S1)**. The resultant particles yielded multiple immunogens with defined percentages of GT5 WT protomers **(Fig. 1A)**. The four variant particles varied in their percent binding activity of germline VRC01 normalized to full eOD-GT5 WT 60mer **(Fig. 1A)**. The normalized percent activities (measured by KinExa) relative to GT5 WT were 79%, 56%, 26%, and 12% **(Table S1)**. Thus, we named our immunogens based on decreasing order of WT GT5 binding activity: eOD-GT5 60mer_100_, eOD-GT5 60mer_79_, eOD-GT5 60mer_56_, eOD-GT5 60mer_26_, eOD-GT5 60mer_12_, eOD-GT5 60mer_0_. For clarification, eOD-GT5 60mer_100_ is normal “WT” eOD-GT5 60mer and eOD-GT5 60mer_0_ is also known as “KO” eOD-GT5 60mer. An alternative description of the mosaic immunogens would be the copies of WT eOD subunits per nanoparticle. Based on decreasing percentage this would be: 60-copies (100%, aka WT), 47-copies (79%), 34-copies (56%), 16-copies (26%), 7-copies (12%), and 0-copies (0%, aka KO). We confirmed mosaicism of these particles by a mixed sandwich ELISA utilzing antibodies specific for the WT or KO only subunits **(Fig. S1A-B**). As expected only mosaic particles were readily detectable in this specific ELISA **(Fig. S1B)**. All mosaic particles bound similarly as expected, due to the counteracting nature of the mixed-epitope sandwhich ELISA. In contrast, the control particles of WT-only, KO-only, and a cocktail of WT-only and KO-only 60mers showed signal below the limit of detection **(Fig. S1B)**. Our mosaic particles were of equal shape morphologically as revealed by negative stain EM **(Fig. 1B)** and size by SEC **(Fig. S1C).** This enabled us to study the impact of epitope valency independent of immunogen size, as antigen size is known to directly affect B cell priming by differential immunogen drainage characteristics.^5^ In sum, these precisely engineered mosaic immunogens enabled us to study the effects of B cell epitope valency in a strictly controlled fashion.

We evaluated various mosaic immunogens for their ability to prime physiologically rare VRC01^gHL^ precursors (1 in 10^6^ precursor frequency) post vaccination **(Fig. 1C)**. Additionally, we included naïve mice in all experiments that have received VRC01^gHL^ B cells as a negative control. We primed separate cohorts of mice with these immunogens (10µg per mouse) in alum. On day 8, resultant mice exhibited a striking stepwise decrease in the VRC01^gHL^ B cell occupancy of GCs that was dependent on B cell epitope valency **(Fig. 1D-E)**. This correlation of VRC01^gHL^ GC B cell response with epitope valency was maintained throughout the peak of the GC reaction in mice **(Fig. 1E)**. The frequency of total GCs was equivalent between all valency groups by flow cytometry **(Fig. S1D)**, highlighting the uniform immunogenicity of the mosaic immunogens. In agreement with this, class switch recombination (CSR) of VRC01^gHL^ GC B cells was also similar between groups throughout all time points **(Fig. S1E)**. The impact of antigen valency on VRC01^gHL^ GC B cell responses was also observed in the absolute number of VRC01^gHL^ GC B cells **(Fig. S1F**). We also evaluated the induction of Tfh by our various mosaic immunogens **(Fig. S1G)**. As expected, Tfh and GC Tfh cell frequency as well as ICOS expression induced by various mosaic immunogens were similar across all groups in both early GC **(Fig. S1H-J)** and peak GC (**Fig. S1K-M**) reactions. We concluded that antigen valency plays a key role in defining the early competitive environment in GCs.

Next, we hypothesized that antigen valency could affect the durability of the GC reaction. To answer this question, we immunized separate cohorts of mice under the same conditions with mosaic eOD-GT5-60mers of varying valency. We then tracked the GC response over one year post prime. We could reliably detect VRC01^gHL^ GC responses at 3- and 6-months post immunization only in mice immunized with eOD-GT5 60mer_56_ mosaic immunogen or higher valency **(Fig. 1F)**. At 9 months post immunization, only a few mice that were immunized with eOD-GT5 60mer_100_ mosaic immunogen had detectable, albeit minor, VRC01^gHL^ GC responses **(Fig. 1F)**. All groups of mice had resolved VRC01^gHL^ GC responses by 12 months post immunization **(Fig. 1G)**. We did note that the percent normalized activity of our mosaic particles correlated with the VRC01^gHL^ GC response across a range of time points with strongest correlations noted early in the response **(Fig. 1H)**. Taken together the data reveals that antigen valency has a critical role not only in defining the early competitive landscape of GCs but also the durability of the GC reaction.

Flow cytometric data does not reveal spatial information. Since GCs exist as microanatomical islands, and GC B cells are not known to transit between individual GCs, clonal competition is largely considered to be localized to individual GCs.^32,69^ Are the effects we see from antigen valency due solely to improved GC seeding by VRC01-class precursor B cells? Or is clonal competition within individual GCs altered? To start to answer such questions, we developed a semi-high throughput histology pipeline (see methods) to quantitate VRC01^gHL^ B cells in GCs. Assessment of GCs on day 8 from mice immunized with mosaic particles ranging from eOD-GT5 60mer_12_ to eOD-GT5 60mer_100_ revealed that total GC size was equivalent **(Fig. S1N)**, reiterating that each mosaic particle was equally immunogenic globally. However, the VRC01^gHL^ GC response in both quantity of GCs containing any VRC01^gHL^ cells, and their relative immunodominance within individual GCs, differed. Interestingly, antigen valency induced more of a thresholding effect in regard to seeding of individual GCs by VRC01^gHL^ B cells. Immunogens with a valency of eOD-GT5 60mer_56_ or greater induced a relatively consistent frequency of GCs containing any VRC01^gHL^ B cells (9-12%). However, lower valency particles eOD-GT5 60mer_12_ or eOD-GT5 60mer_26_ were only detected in 2-4% of GCs **(Fig. 1I-J, Fig. S1O)**. Lastly, we assessed the ability of VRC01-class GCs to undergo clonal bursting. VRC01^gHL^ GC B cells existed in greater quantity within individual GCs when primed with higher valency immunogens on days 8-20 of the GC reaction **(Fig. 1K-L, Fig. S1P)**. Notably, a progressive increase in percent occupancy of individual GCs by VRC01^gHL^ GC B cells was observed from days 8 through 20 only in mice primed with an antigen valency of eOD-GT5 60mer_26_ or greater **(Fig. 1L-M).** While VRC01^gHL^ GC B cells were detectable in mice primed with eOD-GT5 60mer_12_, they never appeared to undergo significant clonal bursting within individual GCs **(Fig. 1M)**. This suggests that antigen valency impacts not only the seeding of individual GCs by VRC01^gHL^ precursor B cells but also the interclonal competition and immunodominance within individual GCs.

### Antigen valency has minimal impacts on somatic hypermutations but impacts clonal diversity

To evaluate if antigen valency affects SHMs in VRC01^gHL^ GC B cells, we performed single cell sorting **(Fig. S2A)** followed by reverse transcription, nested PCR, and sanger sequencing^70^ of heavy and light chains. This approach was chosen because the recovery of these cells was too low within individual groups for our 5’ V(D)J 10x genomics pipeline. We sorted and sequenced approximately 7532 VRC01^gHL^ GC B cells from days 8, 16, and 36 post immunization in mice primed with various valency immunogens **(Fig. 1C, Fig. S2A)**. We found that total heavy and light chain amino acid mutations were similar between mice primed with either eOD-GT5 60mer_100_, eOD-GT5 60mer_79_, eOD-GT5 60mer_56_, or eOD-GT5 60mer_26_ on both early **(Fig. S2B-C, Table S2)** and late time points (**Fig. 2A-B, Table S2).** This similarity was also observed at the nucleotide level **(Fig. S2D-E)**. VRC01^gHL^ GC B cells from mice primed with eOD-GT5 60mer_12_ were too infrequent to sort. Evaluation of individual residue mutations also showed a similar positional pattern across all groups in heavy **(Fig. 2C)** and light chains **(Fig. 2D)** on day 36. This similarity was also noted at earlier time points in the GC reaction for both heavy chains **(Fig. S2F-G)** and light chains **(Fig. S2H-I)**. Assessment of several VRC01-class mutations in heavy chains **(Fig. 2E, Fig. S2J)** and light chains **(Fig. 2F, Fig. S2K)** showed similar levels of mutation across different valency groups on both days 16 and 36. Accrual of key VRC01-class mutations^11^ was similar across groups on day 16 **(Fig. S2L)** and day 36 **(Fig. 2G),** with a small tendency for increased key mutations in mice primed with our highest valency immunogen on day 36. In sum, our analysis revealed that antigen valency does not dramatically affect the accrual of total or individual SHMs.

**Figure 2:**
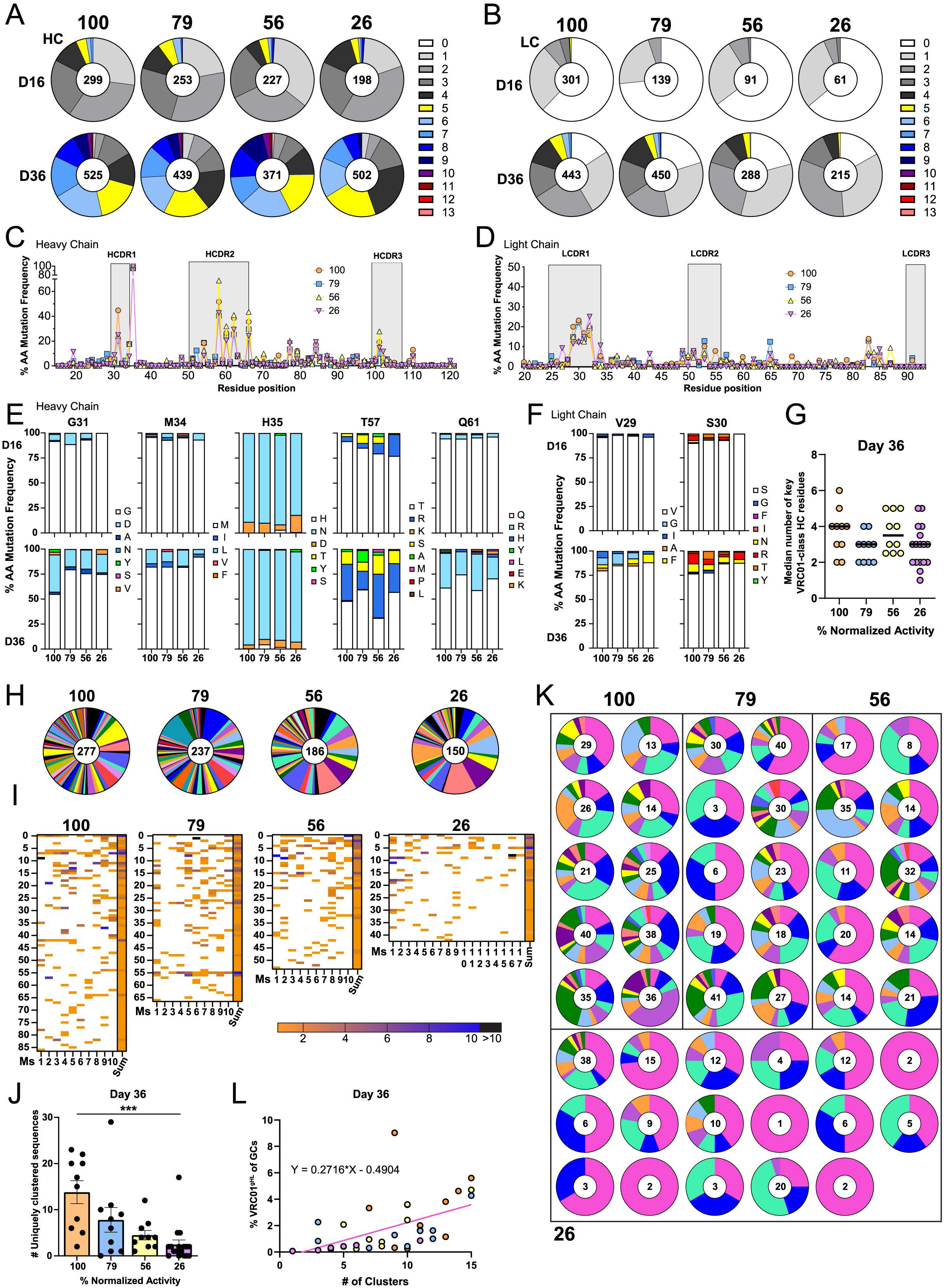
Antigen valency has minimal impacts on somatic hypermutations but impacts clonal diversity. **A-B)** Circle charts represent the fraction of heavy chain (HC) **(A)** and light chain (LC) **(B)** sequences that acquired the indicated number of amino acid (aa) mutations at days 16 and 36 post immunization. Total individual sequences are displayed in circle centers. **C-D)** Frequency of mutation at each residue position within the VRC01^gHL^ **(C)** heavy chain and **(D)** light chain on day 36. **E-F)** Frequency of key VRC01 **(E)** heavy chain and **(F)** light chain mutations at Days 16 and 36. **G)** The 90th percentile values for key VRC01-class heavy chain residues from days 16 and 36. Dots represent individual mice with bars indicating the median of 90th percentile values for each group. **H)** Circle graphs depicting the number of unique clusters by color from the total number of paired clones (shown in circle centers) recovered from mice immunized with mosaic nanoparticles, pooled across experiments. Nanoparticle groups were clustered independently. **I)** Heatmap of relative contribution of each mouse to the clusters depicted in H. Length of heatmap corresponds to absolute number of clusters recovered. **J)** Number of sequences that belonged to a cluster unique to individual mice as shown in I. Mean ± SEM. **K)** Circle graphs depicting the number of unique clusters (by color) from the total number of paired clones recovered (shown in circle centers) from individual mice. Each mouse was clustered independently. **L)** Graph comparing the number of individual clusters calculated per mouse to matched flow cytometry data of the percent VRC01^gHL^ B cells in GCs, as shown in Fig. 1E. Total sequences obtained from mosaic nanoparticle immunizations were pooled across experiments and mice. E = independent experiment. D36 E = 2-5 for 100%, 79%, 56%, and 26%, n = 5-22 mice for HCs & LCs. **G, J)** Kruskal-Wallis with Dunn’s correction. **p<0.01, ***p<0.001. See also Figure S2 and Table S2.

Lastly, we conducted analysis of paired VRC01^gHL^ sequences to evaluate the impacts of antigen valency on clonal diversity on day 36 post immunization. Sequences were grouped by nanoparticle valency group **(Fig. 2H-J)** or further subdivided by individual mice **(Fig. 2K-L).** Sequences were clustered and assessed using Sequencing Analysis and Data Library for Immunoinformatics Exploration (SADIE)^11^ (see methods). We further analyzed these clusters with the Simpson’s Index of Diversity **(Table S3)**, which considers both richness (the number of different clusters present) and evenness (the relative abundance of each cluster), giving more weight to more common clusters and decreasing as one cluster becomes more dominant. We found that by day 36 the diversity of VRC01^gHL^ GC B cells tended to decrease as antigen valency decreased. In mice primed with OD-GT5 60mer_100_, eOD-GT5 60mer_79_, eOD-GT5 60mer_56_, or eOD-GT5 60mer_26_, the mean (S.D.) Simpson’s indices were 0.88(0.04), 0.86(0.10), 0.81(0.12), and 0.64(0.34), respectively **(Table S3, Fig. S2M)**. This trend was seen even when the lowest valency group was oversampled (17 mice assessed vs 10 mice) to account for reduced GC B cell numbers **(Fig. 2H-I)**. Further assessment of unique sequence clusters of VRC01^gHL^ GC B cells revealed that the total number of unique clones present within each mouse was reduced as antigen valency was decreased **(Fig. 2J)**. This was seen when sequences from individual mice were clustered as well **(Fig. 2K)** and appeared to be related to frequency of VRC01^gHL^ B cells in GCs within individual mice **(Fig. 2L)**. This is in line with our observation that the number of VRC01^gHL^ GC B cells able to effectively compete within individual mice was substantially reduced in mice primed with low valency immunogens **(Fig. 1G, Fig. S1F).** In sum, antigen valency affects absolute diversity of BCR sequences that are produced in the GC reaction. In turn, this may impact mutational progression of B cells toward desired neutralizing and protective immune responses.

### Formation and durability of immunological memory is driven by antigen valency

Next, we asked if the production and durability of MBCs and LLPCs were affected by antigen valency. MBCs and LLPCs are key terminal components of productive vaccine immune responses. MBCs are thought to serve as a protective “library” of clones that can quickly differentiate into antibody secreting cells upon re-exposure to homologous or heterologous viral variants.^46,71–73^ MBCs have been shown to last up to a lifetime in humans in highly successful vaccines such as vaccinia.^45^ LLPCs are thought to be the direct cell type responsible for providing durable vaccine immunity over a lifetime.^12,47,48,74^ LLPCs reside in the bone marrow and secrete antibody into the serum for decades in the most successful vaccines.^12,47,48,75^

To ascertain if antigen valency impacted MBC responses, we assessed the formation and durability of VRC01^gHL^ MBCs in mice immunized with various valency immunogens up to one year post prime as in **(Fig. 1C)**. We found that VRC01^gHL^ MBCs formed in a manner dependent on antigen valency **(Fig. 3A-B)**. At one and three months post prime VRC01^gHL^ MBCs were detectable in mice immunized with eOD-GT5 60mer_26_ or higher **(Fig. 3B-C)**. This threshold decreased over time. By one year post prime, only mice primed with medium-valency immunogens (i.e. eOD-GT5 60mer_56_) or higher had detectable VRC01^gHL^ MBCs **(Fig. 3B-C)**. This response was also noted in the absolute number of VRC01^gHL^ MBCs recovered **(Fig. S3A).** Qualitative evaluation of VRC01-class MBCs in the various valency conditions suggested that antigen valency had some modest impacts on driving the type of MBC that forms. VRC01-class MBCs that were likely GC derived (i.e. mostly triple positive for the memory markers CD73, PDL2, and CD80)^76^ appeared to develop slightly later in the GC reaction under lower valency priming conditions **(Fig. S3B)**. Additionally, class-switched VRC01-class MBCs were more likely to be present in mice primed with high valency immunogens at later time points post vaccination **(Fig. S3C)** The VRC01^gHL^ MBC response did correlate with the valency of the immunogen at all time points tested **(Fig. 3D)**. There is substantial interest in defining early predictors of long-term vaccine durability. In this vein, we noted that the peak (day 16) of the early VRC01^gHL^ GC reaction also correlated with later time points of VRC01^gHL^ MBCs **(Fig. 3E)**. Together the data reveals that antigen valency drives the formation and durability of MBC response post vaccination.

**Figure 3:**
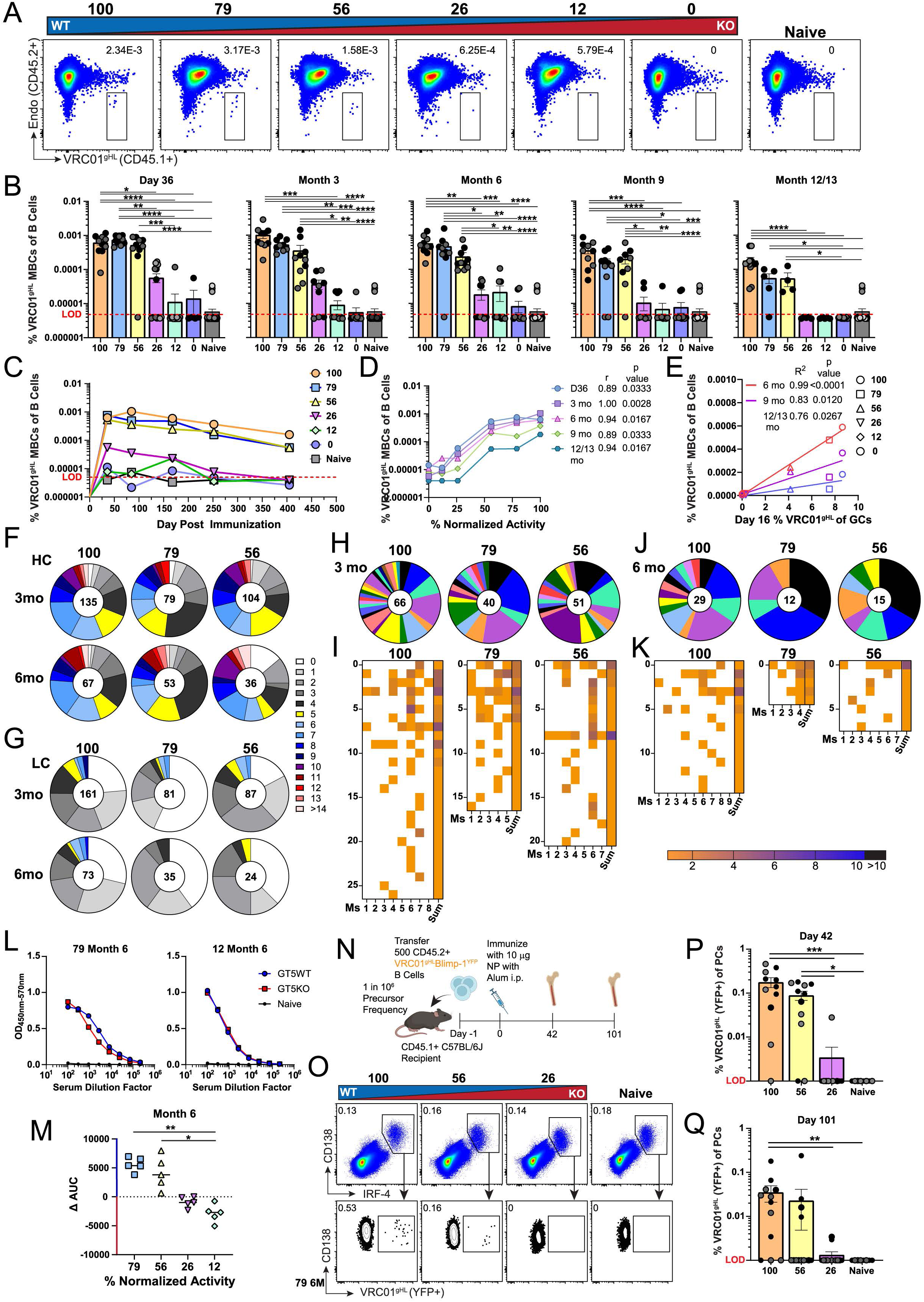
Formation and durability of immunological memory is driven by antigen valency. **A)** Representative flow cytometry from month 6 for quantifying VRC01^gHL^ MBCs from mice immunized with mosaic nanoparticles. Pre-gated on total MBCs SSL/CD4-/B220+/GL7-/CD38+/IgD-/CD138-. **B)** Frequency of VRC01^gHL^ MBCs positive for at least one memory marker (CD73, CD80, or PDL2) among total B cells over time. n= 5 mice/group. **C)** Line graph denoting VRC01^gHL^ frequency in MBCs per group over time as in B. **D)** Spearman coefficient analysis for correlation of mosaic nanoparticle normalized activity and average frequency of VRC01^gHL^ MBCs as in C. **E)** Linear regression analysis comparing average VRC01^gHL^ GC B cell frequencies within GCs from day 16 to average VRC01^gHL^ MBC frequencies among total B cells at months 6, 9 and 12. Best fit line is shown. n= 4-5 mice/group. **F-G)** Circle charts represent the fraction of heavy chain **(F)** and light chain **(G)** sequences from VRC01^gHL^ MBCs that acquired the indicated number of aa mutations at months 3 and 6 post immunization. Total individual sequences are displayed in circle centers. **H)** Circle graphs depicting the number of unique clusters (by color) within the total number of recovered paired clones (shown in circle centers) from 3 months. Nanoparticle groups were clustered independently. **I)** Heatmaps denote the relative contribution from each mouse to each cluster depicted in H at 3 months. Length of heatmap corresponds to absolute number of clusters recovered. **J)** Circle graphs depicting the number of unique clusters (by color) within the total number of recovered paired clones (shown in circle centers) from 6 months. Nanoparticle groups were clustered independently. **K)** Heatmaps denote the relative contribution from each mouse to each cluster depicted in J at 6 months. Length of heatmap corresponds to absolute number of clusters recovered. **L)** Representative month 6 serum IgG ELISAs on mice immunized with 79% (left) and 12% (right) GT5 mosaic nanoparticles against GT5 WT (blue) and GT5 KO (red). **M)** CD4bs WT epitope specific responses as determined by subtracting the KO area under the curve from the WT. Shapes indicate individual mice. n = 5 mice/group. **N)** Schematic of the VRC01^gHL^Blimp-1^YFP^ transfer experiments. **O)** Representative flow cytometry from day 42 for quantifying VRC01^gHL^Blimp-1^YFP^ plasma cells. **P)** Frequency of VRC01^gHL^Blimp-1^YFP^ plasma cells among total plasma cells at day 42 and **Q)** day 101 post prime with mosaic nanoparticles. Bar indicates mean; error bars indicate SEM. E = independent experiment. **B-E)** E = 2; **F-K)** E = 2; **L-M)** E = 1; **P)** E = 2. **B, M, P)** Kruskal-Wallis with Dunn’s correction. *p<0.05, **p<0.01, ***p<0.001, ****p<0.0001. Shades of grey dots indicate independent experiments. See also Figure S3 and Table S2.

We next asked if antigen valency impacted SHMs in VRC01^gHL^ MBCs. To answer this question, we single cell sorted **(Fig. S3D)** and sequenced rare VRC01^gHL^ MBCs from 3- and 6-months post immunization from mice immunized with medium- to high-valency particles (56%-100% or approximately 34-60 copies of the WT subunit). Significant levels of mutation were observed in all groups. The total heavy chain and light chain amino acid mutations were similar between groups at both 3- and 6-months post prime **(Fig. 3F-G, Table S2)**. This was also true at the nucleotide level **(Fig. S3E-F).** Moreover, specific VRC01-class mutations were also similar at both time points in both heavy chains **(Fig. S3G-H)** and light chains **(Fig. S3I-J)**. The total mutational load in VRC01^gHL^ MBCs at any one timepoint was equal or greater than the mutational load detected in VRC01^gHL^ GC B cells on day 36 post immunization **(Table S2)**. The relative diversity of paired sequences was similar in mice primed with medium- or high-valency immunogens, with a tendency for greater diversity in mice primed with the highest valency immunogen at both 3 months **(Fig. 3H-I)** and 6 months **(Fig. 3J-K)** post prime **(Table S3)**. Key VRC01-class mutations were similar in VRC01^gHL^ MBCs recovered from mice primed with medium to high valency immunogens **(Fig. S3K-L)**. Taken together antigen valency has minimal impacts on individual SHMs in MBCs and relative diversity in mice immunized with medium to high valency immunogens. However, we were unable to recover VRC01^gHL^ MBCs from mice primed with immunogens with a lower valency than the eOD-GT5 60mer_56_ group, as these cells were exceedingly rare **(Fig. 3C).** This suggests that multivalency promotes absolute diversity in the MBC compartment through overall competitive expansion of B cells post vaccination.

LLPCs are terminally differentiated, principally reside in specialized bone marrow niches, and are the major drivers of long-lived serum antibody responses.^13,47–49,74^ Our first indication that LLPCs may be impacted by antigen valency was our observation of the epitope-specific antibody response to our mosaic immunogens. Serologically, we assessed CD4bs epitope-specific IgG responses to our various mosaic immunogens by conducting ELISAs for both eOD-GT5 60mer WT and eOD-GT5 60mer KO immunogens. The relative binding to WT versus KO gave us an indication of the CD4bs epitope-specific serum antibody response. The epitope-specific serum antibody response for either WT or KO CD4bs-specific epitopes also tracked with respective epitope valency **(Fig. 3L-M, Fig. S3M-P)**. To directly assess VRC01^gHL^ specific LLPC formation in our model system we crossed our VRC01^gHL^ mice to the Blimp-1^YFP^ reporter mouse,^77^ generating VRC01^gHL^Blimp-1^YFP^ mice **(Fig. 3N)**. This mouse effectively labels approximately 50 percent of LLPCs in bone marrow.^77^ This circumvented the challenge that our tracking allele (CD45.1/2 a.k.a B220) is downregulated on LLPCs. We transferred VRC01^gHL^Blimp-1^YFP^ B cells into CD45.1 mice at a physiologically relevant precursor frequency of 1 in 10^6^ B cells and immunized with a range of mosaic eOD-GT5 nanoparticle immunogens **(Fig. 3N)**. On day 42 post prime, we found that VRC01^gHL^ LLPC formation in the bone marrow was dependent on antigen valency as mice primed with eOD-GT5 60mer_56_ or above had readily detectable VRC01^gHL^ LLPCs **(Fig. 3O-P)**. Moreover, by day 101 there appeared to be a substantial loss of VRC01^gHL^ LLPCs in mice primed with medium-valency immunogens (eOD-GT5 60mer_26_ or lower valency) compared to mice primed with higher valency immunogens (eOD-GT5 60mer_56_ and higher) **(Fig. 3Q)**. This was also observed in the absolute number of VRC01^gHL^ LLPCs **(Fig. S3Q-R)**. In mice primed with our highest valency immunogen, 11 of 12 mice (91.6%) had detectable VRC01^gHL^ LLPCs develop on day 42 post immunization **(Fig. 3P)**. This dropped slightly by day 101 to 9 of 12 mice (75%) having reliably detectable VRC01^gHL^ LLPCs in the BM **(Fig. 3Q)**. Similarly, when mice were primed with eOD-GT5 60mer_56_, 10 of 12 mice (83.3%) had early detectable VRC01^gHL^ LLPCs within the BM on day 42 post immunization **(Fig. 3P)**. However, this dropped precipitously by day 101. On day 101, mice primed with eOD-GT5 60mer_56_ had only 4 of 13 mice (30.7%) with detectable VRC01^gHL^ LLPCs in the BM **(Fig. 3Q)**. In sum, the data suggests that antigen valency promotes both the formation and durability of the LLPC compartment.

### Antigen valency drives memory B cell formation and diversity in a prime-boost regimen

Next, we wondered if the advantage conferred by high valency immunogens in driving productive VRC01^gHL^ B cell responses was because mice received only a priming dose of immunogen. Would this valency advantage disappear if mice received a booster immunization? Most subunit vaccines are given on a vaccine schedule of 0, 1, and 6 months or 0 and 6 months.^75^ Notably, most preclinical murine vaccination models often boost only at the earliest time points possible (e.g. ∼1 month). To investigate whether antigen valency influences the formation of VRC01-class MBCs within a human-like prime/boost regimen, we primed mice containing rare VRC01^gHL^ B cell precursors with eOD-GT5 mosaic immunogens of varying avidities. Then at six months post-prime we boosted each cohort of mice with respective homologous mosaic immunogens **(Fig. 4A)**. At 42 days post boost (day 222 post prime) we found that higher valency immunogens induced a greater magnitude of the VRC01^gHL^ MBC response **(Fig. 4B-C).** CSR among VRC01^gHL^ MBCs was similar across groups with a tendency for increased IgM memory in mice primed and boosted with our lowest valency immunogen, eOD-GT5 60mer_12_ **(Fig. S4A-B)**. The phenotype of VRC01^gHL^ MBCs also varied based on the immunogen valency. VRC01^gHL^ MBCs expressing all three memory markers (CD73, PDL2, CD80), suggesting GC origin, were increasingly prevalent in mice primed and boosted with higher valency immunogens **(Fig. S4C-D)**. A greater phenotypic diversity of VRC01^gHL^ MBCs was also observed in mice primed and boosted with higher valency immunogens **(Fig. S4D)**. To assess the functional impact of homologous boosting with various valency immunogens on the recall activity of VRC01^gHL^ MBCs, we performed epitope-specific IgG ELISAs pre and post boost for the CD4bs WT and CD4bs KO epitopes. As expected, we observed a substantial increase in total antigen antibody titer in all groups **(Fig. 4D)**. Notably, following boosting, the CD4bs WT epitope-specific antibody titer dropped as the valency of the immunogen was reduced **(Fig. 4D-E).** In sum, antigen valency plays a dominant role in driving the magnitude of the MBC response in a homologous prime/boost regimen.

**Figure 4:**
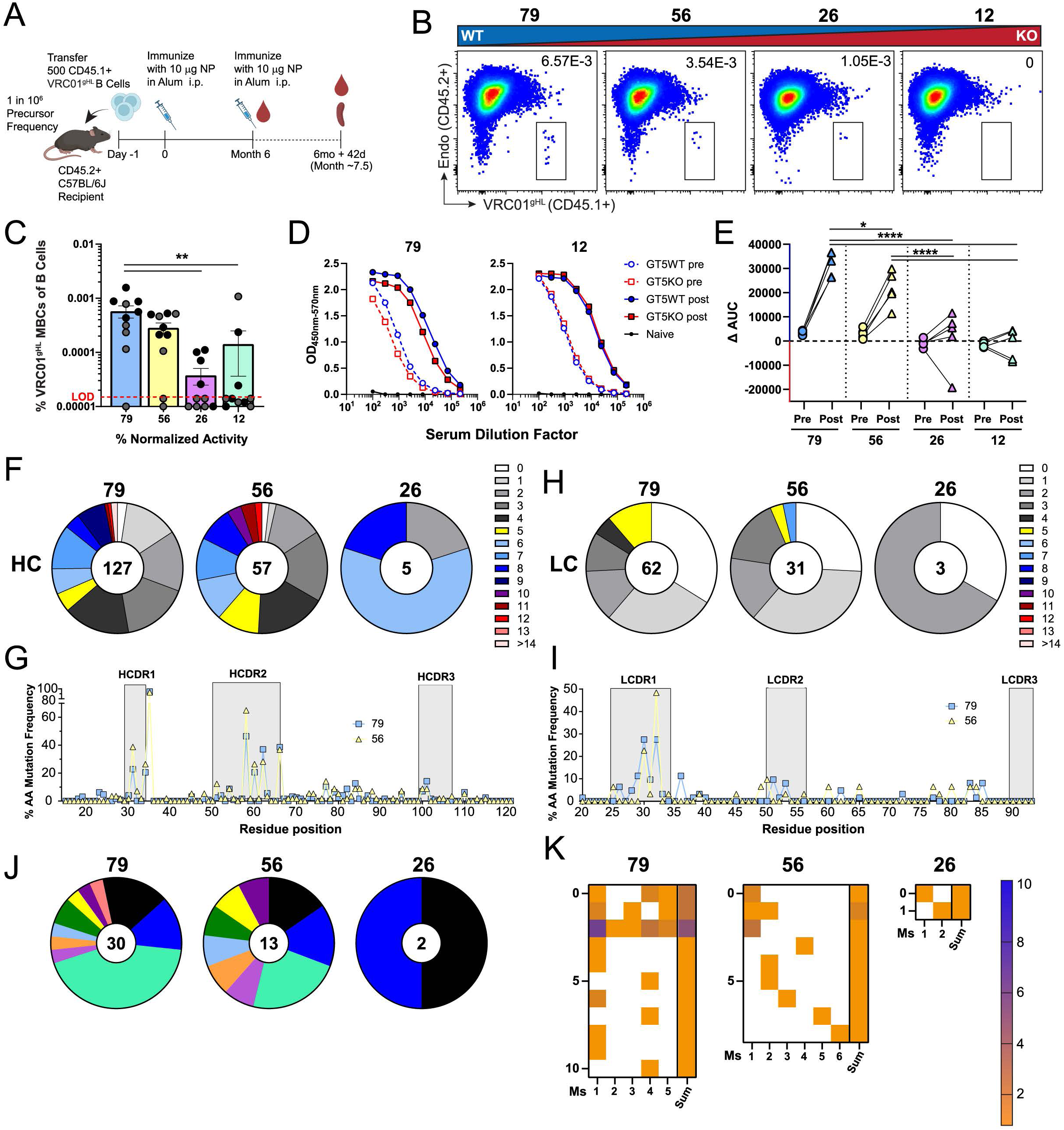
Antigen valency drives memory B cell formation and diversity in a prime-boost regimen. **A)** Schematic of the VRC01^gHL^ prime and boost strategy. **B)** Representative flow cytometry from day 42 post boost for quantifying VRC01^gHL^ MBCs from mice immunized with mosaic nanoparticles. Pre-gated on total MBCs SSL/CD4-/B220+/GL7-/CD38+/IgD-/CD138-. **C)** Frequency of VRC01^gHL^ MBCs among total B cells. n = 5 mice/group. Error bars indicate SEM. **D)** Representative serum IgG ELISAs on mice immunized with 79% (left) and 12% (right) GT5 mosaic nanoparticles from month 6 prior to boosting and day 42 post boost against GT5 WT (blue) and GT5 KO (red). **E)** CD4bs WT epitope specific responses as determined by subtracting the KO area under the curve from the WT from pre and post boost as in D. Dots and triangles indicate individual mice. n = 5 mice/group. **F)** Circle charts representing the fraction of HC sequences from VRC01^gHL^ MBCs that acquired the indicated number of aa mutations at day 42 post boost. Total individual sequences are displayed in the center. **G)** Frequency of mutation at each residue position within VRC01^gHL^ HCs from VRC01^gHL^ MBCs on day 42 post boost. **H)** Circle charts representing the fraction of LC sequences from VRC01^gHL^ MBCs that acquired the indicated number of aa mutations at day 42 post boost. Total individual sequences are displayed in the center. **I)** Frequency of mutation at each residue position within VRC01^gHL^ LCs from VRC01^gHL^ MBCs on day 42 post boost. **J)** Circle graphs depicting the number of unique clusters (by color) within the total number of recovered paired clones (shown in circle centers). Nanoparticle groups were clustered independently. **K)** Heat graphs denote the relative contribution from each mouse to each clonal cluster depicted in J. **C, E)** Kruskal-Wallis with Dunn’s correction. *p<0.05, **p<0.01,****p<0.0001. Shades of grey dots indicate independent experiments. See also Figures 1 and S4 and Table S2.

Next, we sorted **(Fig. S4E)** and sequenced VRC01^gHL^ MBCs on day 42 post boost to ascertain if SHMs were impacted by antigen valency in a prime/boost regimen. Similar amino acid mutational patterns were observed in both heavy **(Fig. 4F-G)** and light chains **(Fig. 4H-I)** with only a few sporadic clones recovered in mice primed with the lowest valency immunogen. This was also observed at the nucleotide level **(Fig. S4F-H)**. Individual VRC01-type mutations were similar across groups in both heavy **(Fig. S4I)** and light chains **(Fig. S4J)**. Evaluation of key VRC01-class heavy chain residue positions was also similar but showed a trend for an improved response in mice primed with higher valency immunogens **(Fig. S4K)**. Paired sequence diversity of VRC01^gHL^ MBCs was reduced as antigen valency was reduced **(Fig. 4J-K, Table S3)**. This appeared to be related to absolute numbers and attributed to the extreme sparsity of VRC01^gHL^ MBCs recovered in our lowest valency group **(Fig. 4J-K)**. In sum, antigen valency has a minimal role in driving individual SHMs but plays a key role in driving the magnitude, durability, and diversity of the MBC responses in a realistic prime-boost regimen.

### Antigen valency regulates interclonal competition within germinal centers

Making use of the VRC01^gHL^ system to study B cell responses to precisely characterized immunogens has substantial advantages including defined precursor frequencies and affinities that are matched to the human repertoire. However, all models have limitations. One inherent limitation of this system is the monoclonal nature of it. To further expand our understanding of the impacts of antigen valency on B cell responses post-vaccination to complex antigens, we deeply interrogated the polyclonal endogenous GC B cell response to our eOD-GT5 60mer mosaic immunogens in mice containing VRC01^gHL^ B cells **(Fig. 1C)**. eOD-GT5 60mers_100_ have been shown to elicit a highly diverse endogenous GC B cell response.^8^ This high level of interclonal competition is more reflective of real-world vaccines than many simple model immunogens that have been utilized to study antigen valency.^78^ Moreover, studying the endogenous GC B cell response to specific eOD-GT epitopes enables us to evaluate the impact of antigen valency in a polyclonal system with high levels of interclonal competition and diverse precursor frequencies and affinities.

Use of avi-tagged, eOD-GT5 specific, flow cytometric probes for WT and KO subunits allowed us to track the endogenous polyclonal epitope specific GC B cell response **(Fig. 5A)**. These endogenous GC B cells were competing against transferred VRC01^gHL^ GC B cells **(Fig. 1C**). The flow probes allowed us to delineate B cells into four distinct populations: CD4bs WT (eOD-GT5 WT^+^KO^-^) (blue box), CD4bs KO (eOD-GT5 WT^-^KO^+^) (red box), double positive (eOD-GT5 WT^+^KO^+^ (shared epitopes outside of CD4bs)), and double negative (eOD-GT5 WT^-^ KO^-^) **(Fig. 5A)**. Interestingly, the endogenous GC B cell response to the CD4bs WT epitope decreased in a stepwise manner with decreasing antigen epitope valency throughout the primary GC reaction **(Fig. 5A-B).** This response resolved by 3 months post immunization **(Fig. 5B)**. The relative GC response to this epitope was correlative with CD4bs WT epitope valency during the primary GC reaction through day 36 **(Fig. 5C**). Notably, this observation was not restricted to CD4bs WT specific GC B cells. The CD4bs KO epitope-specific GC B cell frequencies increased as CD4bs KO^-^epitope valency was increased on the mosaic immunogens **(Fig. 5D-E).** This observation was even detectable at 3 months post immunization **(Fig. 5D)**. This suggests the impact of valency on driving polyclonal competition is likely broadly applicable, regardless of different B cell repertoires responding to a specified complex epitope. CSR was similar in total endogenous GC B cells across groups throughout the GC reaction **(Fig. S5A)**. As expected, the frequency of GC B cells responding to shared epitopes (WT^+^KO^+^) did not vary between the different mosaic immunogens **(Fig. S5B)** and did not correlate with epitope valency **(Fig. S5C),** serving as an internal control. Similarity between valency groups was also observed for the double negative GC compartment **(Fig. S5D-E)**. Together the data reveals that epitope valency plays dominant roles in driving epitope specific GC responses in the context of a complex immunogen with competing epitopes. Moreover, the data suggests that antigen valency plays a role in epitope immunodominance and the durability of epitope-specific GC B cell responses in a polyclonal system.

**Figure 5:**
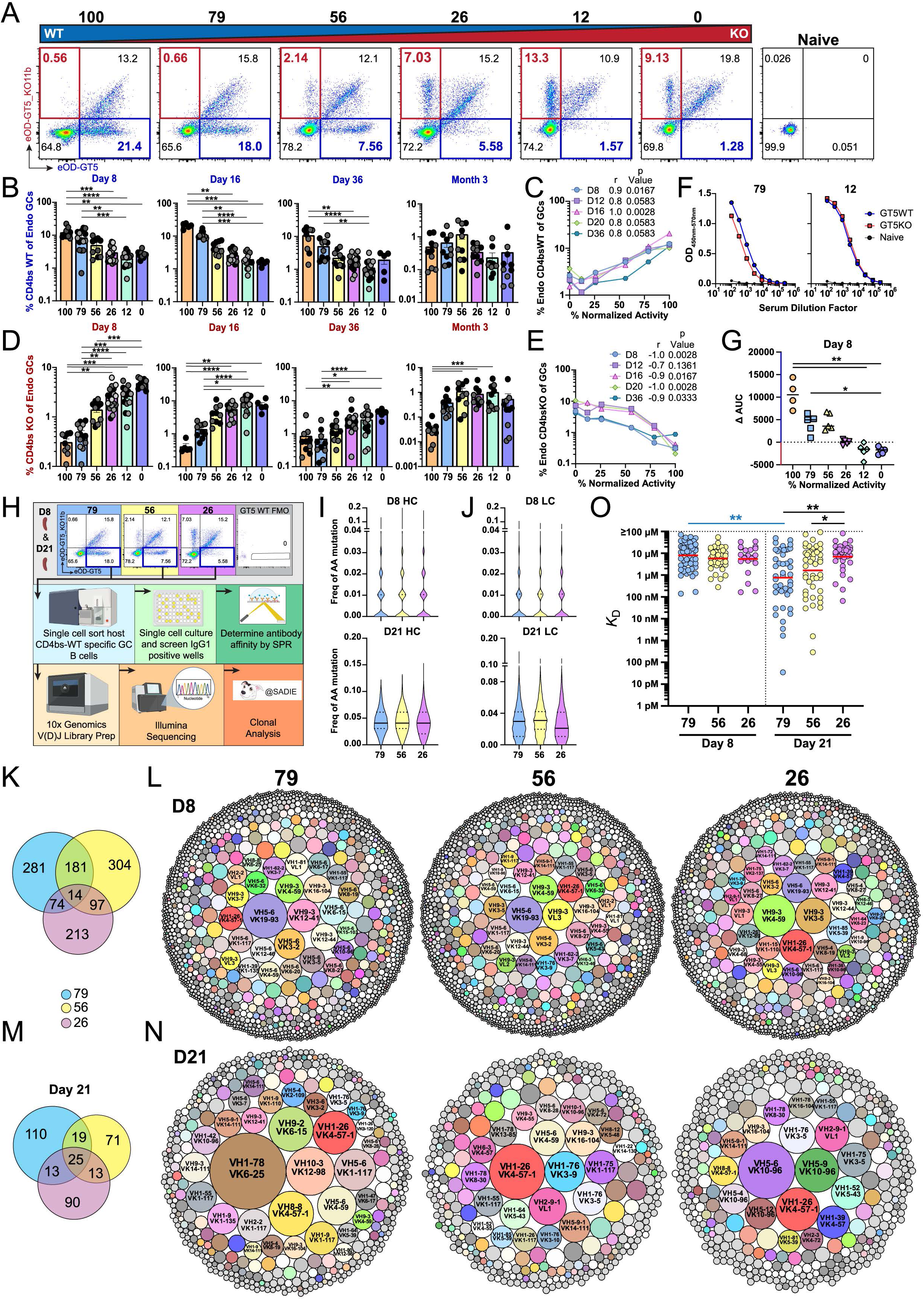
Antigen valency regulates interclonal competition within germinal centers. **A)** Representative flow cytometry from day 16 for quantifying endogenous GC B cell (CD45.2+) binding to eOD-GT5 WT (blue) and KO (red) probes from mice immunized with mosaic nanoparticles. Pre-gated on endogenous total GCs SSL/CD4-/B220+/CD38-/GL7+/CD45.1-/CD45.2+. **B)** Frequency of endogenous GC B cells binding the WT CD4bs. n= 5 mice/group. **C)** Spearman coefficient analysis for correlation between mosaic nanoparticle normalized activity and average endogenous GC binding to the WT CD4bs as shown in B. **D)** Frequency of endogenous GC B cells binding the KO CD4bs. n= 5 mice/group. **E)** Spearman coefficient analysis for correlation between mosaic nanoparticle normalized activity and average endogenous GC binding to the KO CD4bs as shown in D. **F)** Representative day 8 serum IgG ELISAs on mice immunized with 79% (left) and 12% (right) GT5 mosaic nanoparticles against GT5 WT (blue) and GT5 KO (red). **G)** CD4bs WT epitope specific responses as determined by subtracting the KO area under the curve from the WT. Shapes indicate individual mice. n= 4-5 mice/group. **H)** Graphical depiction of workflow for analysis of the endogenous response to mosaic nanoparticles. **I-J)** Frequency of aa mutations in **(I)** HCs and **(J)** LCs of endogenous GC B cells at days 8 and 21. n= 8-9 mice/group. **K)** Venn diagram illustrating the number of unweighted shared and unique clonal families seen more than once from K. n= 8-9 mice/group. **L)** Bubble graph depictions of clonal family abundance in mice immunized with mosaic nanoparticles at day 8. n= 8-9 mice/group. **M)** Venn diagram illustrating the number of unweighted shared and unique clonal families seen more than once from M. n= 8-9 mice/group. **N)** Bubble graph depictions of clonal family abundance in mice immunized with mosaic nanoparticles at day 21. n= 8-9 mice/group. **O)** Quantification of affinity of antibodies derived from cultured CD4bs WT binding GC B cells. **A-E)** E= 1-5; **F-G)** E= 1; **H-N)** E= 2; **O)** E= 1. **B, D)** Kruskal-Wallis with Dunn’s correction. Mean ± SEM. Shades of grey dots indicate independent experiments. **G)** Kruskal-Wallis with Dunn’s correction. **O)** Black bars: Kruskal-Wallis with Dunn’s correction within timepoint. Blue bar: Mann Whitney-U. *p<0.05, **p<0.01, ***p<0.001, ****p<0.0001. See also Figures 1 and S5 and Tables S4-S5.

We found that the CD4bs epitope-specific serum IgG response tracked with CD4bs epitope valency on both days 8 **(Fig. 5F-G)** and 36 **(Fig. S5F-G)**. We also evaluated the serological response to the LS core nanoparticle as a control since this is an equally shared component of all mosaic immunogens. Moreover, since LS is an internal component of the mosaic immunogens, we reasoned that we could use the LS antibody response as a proxy to estimate the *in vivo* stability of our mosaic nanoparticles. The mosaic particles all appeared to elicit equally minimal IgG responses to the LS core **(Fig. S5H-I)**, suggesting they are equally stable *in vivo*. In sum, the early serological response to our mosaic immunogens is dependent on antigen valency.

To assess the qualitative impacts of antigen valency on GC reactions, we next sorted **(Fig. S5J)** and sequenced the endogenous CD4bs WT (eOD-GT5 WT^+^KO^-^) GC population utilizing our 10x genomics 5’ V(D)J pipeline **(Fig. 5H)**. We evaluated GC B cells from mice primed with immunogens of different relative valency. We used “high” (eOD-GT5 60mer_79_), “medium” (eOD-GT5 60mer_56_), and “low” (eOD-GT5 60mer_26_) valency immunogens on days 8 and 21 post immunization **(Fig. 5H)**. We recovered thousands of unique paired clones on day 8 post immunization in each group. SHMs were similar between all groups in heavy and light chains on days 8 and 21 post immunization at the amino acid level **(Fig. 5I-J)** and nucleotide level **(Fig. S5K)**. This was also true in absolute numbers of mutations at the amino acid level **(Fig. S5L)** and nucleotide level **(Fig. S5M)**. CSR was also similar between groups on both days 8 and 21 showing a progression over time toward IgG1 isotype as expected **(Fig. S5N).** The total relative diversity of BCR sequences trended higher with higher valency groups day 8, although the differences appeared to resolve by day 21 **(Fig. S5O, Tables S4)**. Absolute diversity was highest in the higher valency group **(Fig. S5P-Q)**. This was evident as there were almost 3-fold more CD4bs WT epitope-specific GC B cells present in mice primed with eOD-GT5 60mer_79_ when compared with eOD-GT5 60mer_26_ **(Fig. 5A-B).** In essence, antigen valency had minimal impacts on CSR, SHMs, and relative diversity in the polyclonal B cell response, but promoted absolute diversity through competitive expansion of epitope-specific GC B cells.

Next, we assessed if antigen avidity impacted immunodominance patterns at the clonal level. We observed that approximately 70% of total BCR clonal families used in GCs on day 8 post immunization were distinct between groups **(Fig. 5K, Fig. S5R)**. However, when restricted to the top 20 immunodominant clonal families, the majority were shared between groups on day 8 **(Fig. 5L, Fig. S5S-T)**. This suggests that a significant fraction of unique B cells were recruited to GCs based on avidity. Curiously, substantial divergence of clonal usage, and dominance, in clonal families occurred by day 21 post immunization. There was less overlap in the total number of shared clonal families **(Fig. 5M, Fig. S5R).** Assessment of the top 20 immunodominant clonal families between groups showed even further divergence by day 21 between groups when compared to day 8 **(Fig. 5N, Fig. S5U-V).** Overall, the data reveals that antigen avidity plays a substantial role in determining the immunodominance patterns and clonal competition within polyclonal epitope-specific GC responses to a complex immunogen.

We also assessed the qualitative impacts of antigen valency in driving B cell recruitment and affinity maturation in GCs. Using the Nojima single GC B cell culture system,^79,80^ we single cell sorted and expanded over 1000 CD4bs WT epitope-specific endogenous GC B cells from mice primed with high, medium, or low valency immunogens on both days 8 and 21 post immunization **(Fig. 5H)**. This allowed for semi-high throughput population level analysis of 1014 monoclonal antibodies (mAbs) using surface plasmon resonance (SPR) **(Table S5)**. After cell sorting and culture, all mAbs were screened by SPR. Only mAbs that had detectable binding (<100µM K_D_) and were specific for only the CD4bs WT^+^KO^-^ epitope were analyzed, leaving 247 mAbs for analysis. The affinities of CD4bs WT-specific B cells recruited to GCs under various valency conditions were relatively similar on day 8 post-immunization, with geometric mean *K*_D_ values for all mAbs ranging from ∼6µM to ∼8µM **(Fig. 5O)**. By day 21, affinity maturation was most evident in the high-valency group, with a geometric mean *K*_D_ of 0.85µM, representing a 9.44-fold increase **(Fig. 5O, Table S5)**. In contrast, mice primed with medium-valency nanoparticles exhibited more modest changes in affinity, with *K*_D_ values of 1.64µM, corresponding to a 3.51-fold change **(Fig. 5O, Table S5).** Mice primed with low-valency nanoparticles did not undergo appreciable affinity maturation in this timeframe with similar affinities noted on day 8 and 21 (6.33µM and 6.98µM, respectively) representing a 0.91-fold change **(Fig. 5O, Table S5)**. Thus, our data demonstrates that antigen valency had a minimal impact on the affinity of the epitope-binding B cells in the initial recruitment into germinal centers. However, antigen valency promoted markedly accelerated affinity maturation kinetics by day 21 in the highest valency group when compared to the lowest valency group **(Fig. 5O)**. These findings suggest that antigen valency plays a role in shaping the competitive landscape and affinity maturation process within germinal centers.

### The effects of antigen valency are dependent on interconal competition

Next, we sought to investigate mechanisms by which antigen valency impacted B cell priming to enter, compete within, and exit GC reactions. We found that the dominant impacts of antigen valency in driving VRC01-class GC B cell responses and formation of memory were not adjuvant restricted. Immunization of different cohorts of mice with soluble adjuvants RIBI or AddaS03 gave similar results with varying valency immunogens compared to Alum primed mice **(Fig. S6A-B)**. We also wanted to evaluate if our observed decreases in VRC01^gHL^ response with our lower valency mosaic nanoparticles was simply due to the reduced dose of the WT epitope. To formally test this, we compared the ability of VRC01^gHL^ B cells to compete within GCs in mice in one of five conditions, all in alum. These conditions included: (i) 10μg of eOD-GT5 60mer_100_, (ii) a mixture of 9μg:1μg (eOD-GT5 60mer_100_ to eOD-GT5 60mer_0_), (iii) a mixture of 1μg:9μg (eOD-GT5 60mer_100_ to eOD-GT5 60mer_0_), (iv) 1μg of eOD-GT5 60mer_100_, and (v) 10μg of eOD-GT5 60mer_12_. **(Fig. S6C-D)**. The lower dose in condition (iv) was chosen to mimic the epitope dose of the WT subunit on our WT GT5-60mer_12_ mosaic nanoparticle, as 1μg of GT5-60mer_100_ is approximately equal to the WT dose contained within the eOD-GT5 60mer_12_ mosaic nanoparticle. As expected, all groups with a 10μg total dose gave similar total GC resposnes, whereas the 1μg eOD-GT5 60mer_100_ group was comparatively lower **(Fig. S6C)**. Mice primed with either 1μg of eOD-GT5 60mer_100_ or the mixture of 1μg:9μg (eOD-GT5 60mer_100_ to eOD-GT5 60mer_0_) showed an order of magnitude greater response of VRC01^gHL^ GC B cells when compared to mice primed with 10μg of the mosaic WT GT5-60mer_12_ immunogen **(Fig. S6D)**. We interpret this to mean that the effects of mosaic nanoparticles on driving durable vaccine immune responses is not principally due to the increased WT epitope dose in our high valency immunogens. Rather, the data suggests that the multivalent structure of the antigen, and how it directly binds to BCRs on the surface of B cells *in vivo*, plays a dominant role in regulating successful and durable vaccine immune responses.

Next, we tested whether the differential priming of VRC01^gHL^ B cells by high and low valency immunogens to enter, and compete within GCs was dependent on interclonal competition. We hypothesized that interclonal competition could play a role in determining the competitive success of VRC01^gHL^ B cells within GCs. In our mosaic nanoparticles, when the CD4bs WT GT5 epitope decreases, the off-target CD4bs KO epitope increases **(Fig. 1A)**, thus mimicking natural valency differences to subdominant and immunodominant target epitopes on complex viral targets. Experimentally, we addressed this by transferring VRC01^gHL^ B cells at the same precursor frequency (1 in 10^6^ B cells) into mice with a significantly restricted BCR repertoire (MD4 mice) **(Fig. 6A)**. MD4 mice have an efficient transgenic BCR yielding >99% of B cells to be specific for an irrelevant antigen (hen egg lysozyme) and have structurally normal lymphoid tissues **(Fig. S6E)**.^81^ This effectively attenuates vaccine specific endogenous B cell competition. Transfer of B cells into this mouse enabled us to study the impact of interclonal competition on the effect of antigen valency on priming VRC01^gHL^ B cell responses. Total GC responses were similar between groups at time points analyzed **(Fig. S6F)**. We found that low valency eOD-GT5 60mer_26_ nanoparticles were able to prime rare VRC01^gHL^ B cells to enter and compete within GCs in MD4 mice **(Fig. 6B-C)**. CSR of VRC01^gHL^ GC B cells was similar across all groups **(Fig. S6G)**. Moreover, low valency eOD-GT5 60mer_26_ particles were able to drive early VRC01^gHL^ MBC formation at similar levels as high valency eOD-GT5 60mer_100_ immunogens **(Fig. 6D-E).** Phenotypic analysis of VRC01^gHL^ MBC showed similar memory marker usage and levels of CSR across all groups **(Fig. S6H-I)**. In sum, when VRC01^gHL^ B cells were primed with various valency immunogens under a restricted interclonal competition setting, the competitive advantage of VRC01^gHL^ B cells primed with high valency immunogens to enter and compete within GCs disappeared. The data indicates that interclonal competition plays a crucial role in defining the valency thresholds that determine how B cells compete to enter, compete within, and exit GCs as memory post vaccination.

**Figure 6:**
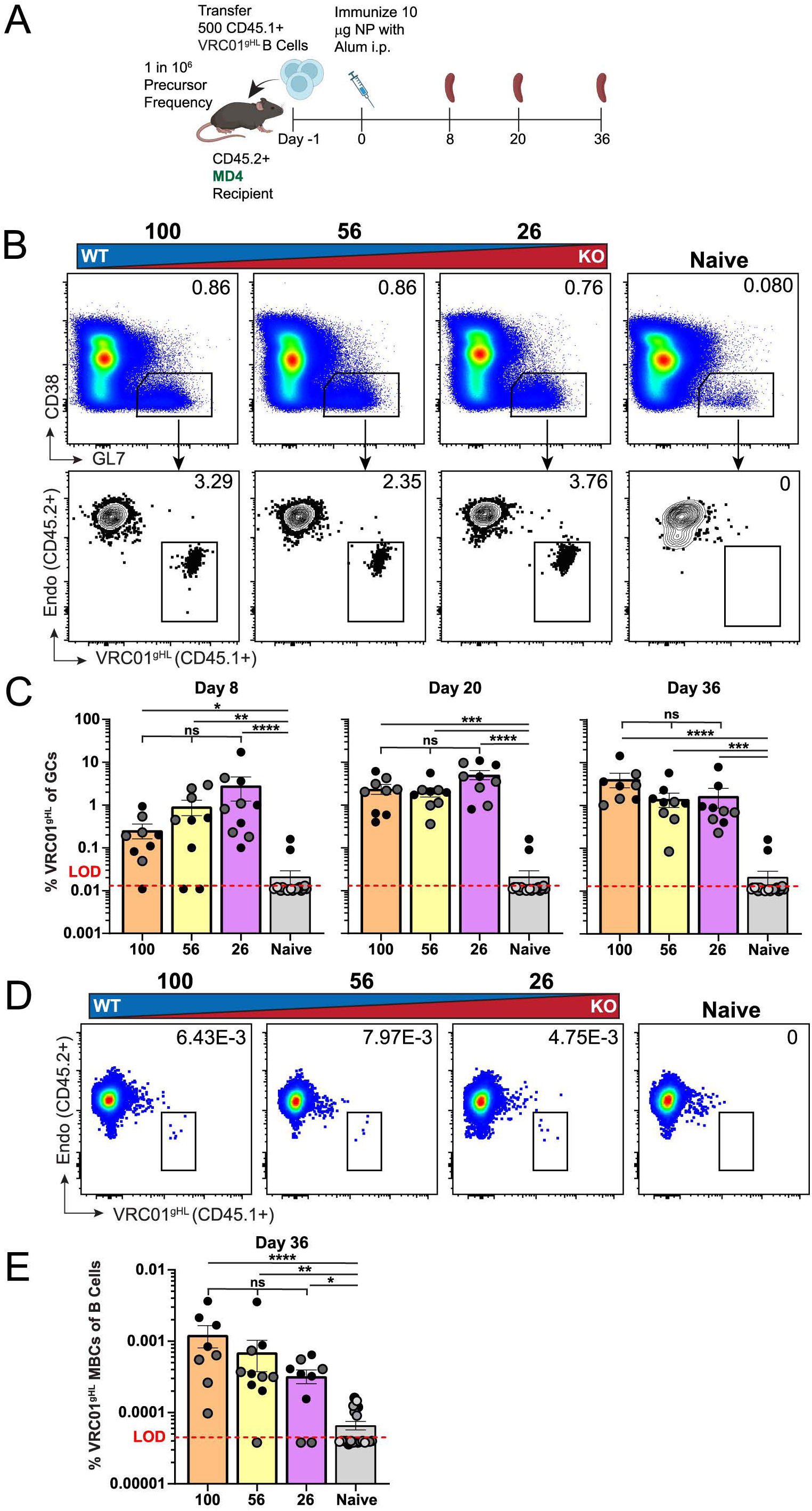
The effects of antigen valency are dependent on interconal competition. **A)** Schematic illustrating the VRC01^gHL^ B cell transfer into a mouse expressing irrelevant antigen receptor (MD4). **B)** Representative flow cytometry from day 20 for quantifying VRC01^gHL^ B cells within total GCs. Pre-gated on SSL/CD4-/B220+. **C)** Frequency of VRC01^gHL^ B cells within total GCs. **D)** Representative flow cytometry from day 36 for quantifying VRC01^gHL^ MBCs. Pre-gated SSL/CD4-/B220+/GL7-/CD38+/IgD-/CD138-. **E)** Frequency of VRC01^gHL^ MBCs positive for at least one MBC marker (CD73, CD80, or PDL2) among total B cells. **C, E)** Kruskal-Wallis with Dunn’s correction. Mean ± SEM. *p<0.05, **p<0.01, ***p<0.001, ****p<0.0001. Shades of grey dots indicate independent experiments. See also Figure 1 and S6.

### Antigen valency drives germinal center and memory B cell responses in authentic human germline models across a range of affinities

Lastly, we asked if our observations regarding the effects of antigen valency in driving successful humoral immune responses in our VRC01^gHL^ mouse model would be reproducible in other models. To test this, we made use of two more stringent next-generation HuGL B cell transfer mouse models. These models utilize authentic human germline VRC01-class BCR sequences that were recovered from healthy human donors.^65^ Moreover, these two models span a 10-fold affinity range. This allowed us to glean insights into the relative contribution of the individual components of antigen avidity, valency and affinity, in driving successful B cell responses post vaccination. In order to test the effects of valency in these models, we produced a new set of mosaic particles using the clinical trial-validated immunogen eOD-GT8 60mer^10,11,28,64^ **(Fig. 7A)**. As with our previous mosaic particles, these immunogens were of equal appearance morphologically by negative stain EM **(Fig. 7A)** and size by SEC (**Fig. S7A).** Furthermore, we also confirmed mosaicism by our mixed-epitope sandwich ELISA (**Fig. S7B)**. These new mosaic particles had slightly different mixed avidities when compared to our GT5-based mosaic particles. We were able to test five independent B cell epitope valency conditions: eOD-GT8 60mer_100_, eOD-GT8 60mer_90_, eOD-GT8 60mer_37_, eOD-GT8 60mer_14_, or eOD-GT8 60mer_10_ (ranging from approximately 60 to 6 copies of WT eOD-GT8) **(Fig. 7A, Table S6)**. As with our previous particles, these mosaic immunogens were equally immunogenic as they induced equivalent total GC responses **(Fig. S7C-D),** as well as levels of CSR (**Fig. S7E-F)**. Moreover, as expected, the induction of Tfh **(Fig. S7G)** and GC Tfh cells **(Fig. S7H)** was equivalent across all mosaic groups.

**Figure 7:**
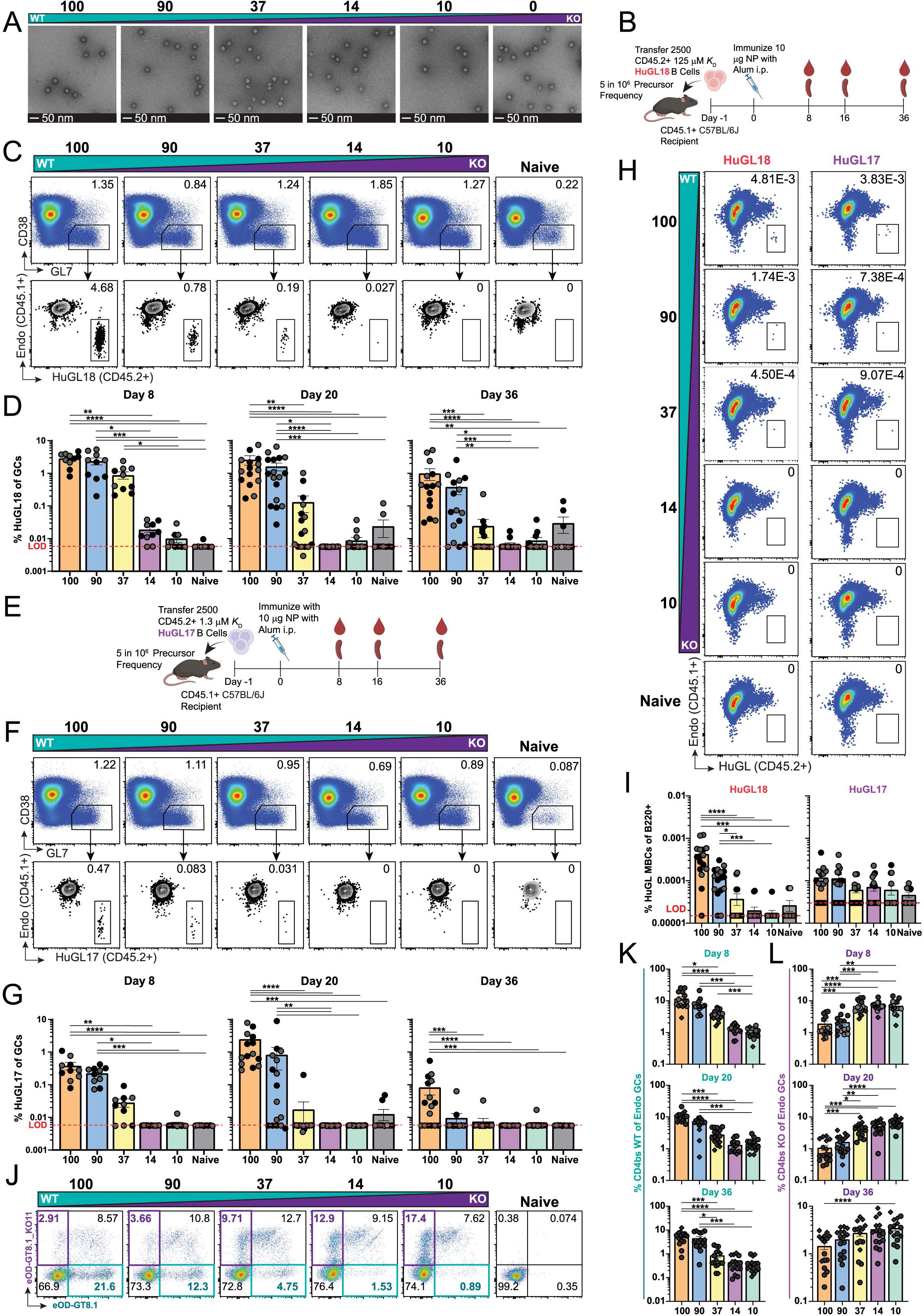
Antigen valency drives germinal center and memory B cell responses in authentic human germline models across a range of affinities. **A)** Negative stain EM images of mosaic nanoparticles that were developed with varying ratios of GT8 WT to GT8 KO. **B)** Schematic of the HuGL18 B cell transfer experiments. **C)** Representative flow cytometry from day 8 for quantifying HuGL18 frequency (CD45.2+) within GCs (CD38-/GL7+). Pre-gated on SSL/CD4-/B220+. **D)** Frequency of HuGL18 B cells within GCs as shown in C. **E)** Schematic of the HuGL17 B cell transfer experiments. **F)** Representative flow cytometry from day 8 for quantifying HuGL17 frequency within GCs (CD38-/GL7+). Pre-gated on SSL/CD4-/B220+. **G)** Frequency of HuGL17 cells within GCs as shown in F. **H)** Representative flow cytometry from day 36 for quantifying HuGL18 and HuGL17 (CD45.1+) MBCs among total B cells. Pre-gated SSL/CD4-/B220+/GL7-/CD38+/IgD-/CD138-. **I)** Frequency of (left) HuGL18 and (right) HuGL17 MBCs positive for at least one MBC marker (CD73, CD80, or PDL2) among total B cells. **J)** Representative flow cytometry from day 20 for quantifying endogenous GC B cell (CD45.2+) binding to eOD-GT8 WT (teal) and KO (purple) probes from mice immunized with GT8 mosaic nanoparticles. Pre-gated on endogenous total GCs SSL/CD4-/B220+/CD38-/GL7+/CD45.1+/CD45.1+. **K-L)** Quantification of the percent of endogenous GC B cells binding the **(K)** WT CD4bs (teal) and **(L)** KO (purple). Circles indicate HuGL18 and diamonds indicate HuGL17. **D, G, I, K-L)** Kruskal-Wallis with Dunn’s correction. Mean ± SEM. *p<0.05, **p<0.01, ***p<0.001, ****p<0.0001. Shades of grey indicate independent experiments. See also Figure S7 and Table S6.

We first tested the impacts of antigen valency on priming our high affinity HuGL18 (∼125nM *K*_D_) B cells starting from physiologically relevant precursor frequencies (5 in 10^6^ precursor frequency) **(Fig. 7B)**. We found that antigen valency played dominant roles in driving seeding and durability of the HuGL18 GC response **(Fig. 7C-D)**. On day 8 post immunization, mice primed with eOD-GT8 60mer_37_ or above were readily detectable within GCs. However, the HuGL18 GC response in mice primed with eOD-GT8 60mer_37_ waned substantially more quickly than mice primed with higher valency immunogens **(Fig. 7D, Fig. S7I)**. We next tested the impacts of antigen valency on priming the medium affinity HuGL17 (1.3µM *K*_D_) VRC01-class B cells to enter and compete within GCs under the same conditions we assessed with HuGL18 B cells **(Fig. 7E)**. Indeed, we found that antigen valency had dominant impacts on driving medium affinity HuGL17 B cells to enter and compete within GCs **(Fig. 7F-G)**. In direct comparison, HuGL17 B cells competed less effectively than the higher affinity HuGL18 B cells **(Fig. S7J)**. Moreover, the difference between medium affinity HuGL17 and high affinity HuGL18 GC B cell responses was exaggerated as antigen valency was reduced **(Fig. S7J)**. This highlights the interdependence of these two variables and the importance of overall antigen avidity in driving productive B cell responses. This agrees with previous observations.^6,8,65^ We also evaluated the impacts of antigen valency on driving MBC formation on day 36 post immunization in both HuGL18 and HuGL17 models **(Fig. 7H)**. We found that antigen valency had stark impacts on the formation of MBCs in both models **(Fig. 7H-I)**. Only HuGL18 MBCs were readily detectable in the highest valency groups **(Fig. 7I)**. Isotype switching of HuGL18 and HuGL17 MBC was not readily affected by antigen valencies in which cells were reliably detectable **(Fig. S7K)**. The impact of antigen valency on phenotype of HuGL MBCs was variable. We generally observed that higher valency immunogens promoted increased levels of CSR and formation of GC derived, triple positive, MBCs **(Fig. S7K-L)**. In sum, antigen valency plays a dominant, but still interdependent role with antigen affinity, in determining successful priming of high and medium affinity B cell precursors to enter, compete within, and exit GC reactions.

The endogenous epitope-specific polyclonal GC B cell response to our eOD-GT8 mosaic nanoparticles was also dependent on antigen valency. There were significantly fewer CD4bs WT (eOD-GT8 WT^+^KO^-^) epitope specific B cells when mice were primed with low valency immunogens **(Fig. 7J-K**). Moreover, this was not unique to the CD4bs epitope as the same dependence on antigen valency was reciprocally observed in the KO epitope (eOD-GT8 WT^-^KO^+^) **(Fig. 7J, L)**. As expected, the endogenous GC B cell response to shared epitopes did not change between groups **(Fig. S7M-N)**. The CD4bs epitope-specific serum antibody response also tracked with antigen valency in mice receiving either HuGL18 **(Fig. S7O-P)** or HuGL17 transferred cells **(Fig. S7Q-R)**. In sum, antigen valency plays a substantial role in promoting productive B cell responses. It also reiterates the complementary and interdependent nature of antigen affinity and valency in driving productive B cell responses post vaccination with clear dominant impacts of antigen valency when starting from physiological precursor frequencies and affinities. Multivalency promotes the entrance, competition within, and exit from GCs to MBC across a range of physiological affinities in authentic human germline B cell models.

## Discussion

Taken together, our study points to the importance of antigen avidity in driving durable and diverse humoral immune responses post vaccination. Specifically, antigen multivalency plays a dominant role in this process. This was true under physiological precursor frequency and affinity conditions, whilst utilizing real-world complex immunogens that elicit high levels of interclonal competition. We found that antigen multivalency promoted the seeding and durability of GCs. Furthermore, antigen multivalency was a key regulator of immunodominance and interclonal competition within GCs. In terms of key long-term vaccine outputs, antigen multivalency promoted the development, diversity, and durability of MBC responses out to over one year post prime. This effect was also seen in a prime boost regimen. Moreover, the generation and durability of epitope specific serum antibody responses as well as LLPC responses in the bone marrow were dependent on multivalency. Mechanistically, the ability of antigen multivalency to drive productive B cell responses was dependent on interclonal competition. Furthermore, our general observations regarding the impacts of antigen multivalency in driving successful B cell responses post vaccination were consistent across a total of five model systems. This included tracking three independent anti-HIV V(D)J humanized knock-in B cells with defined precursor frequencies and affinities, as well as two different endogenous B cell responses to two different sets of mosaic immunogens of varying valencies. The affinity range of B cells tested was wide, ranging from our highest affinity monoclonal HuGL18 B cells (125nM *K*_D_) to the lowest affinity polyclonal B cells responding to the CD4bs KO epitope on eOD-GT5 in mice primed with the eOD-GT5 60mer_79_ (12µM *K*_D_ median) **(Fig. 5O)** representing a 96-fold range. While antigen affinity did certainly act in concert with valency in potentiating B cell responses **(Fig. 7, Fig. S7J)**, antigen multivalency appeared to play a dominant role in driving productive B cell responses.

Our mosaic nanoparticle system contains differential levels of dominant and subdominant epitopes based on the particle that was used. As the WT subunit decreased, the KO subunit increased **(Fig. 1A-B**, **Fig. 7A)**. We found that regardless of epitope, the multivalency, or relative epitope density, was critical in promoting the competitive nature of epitope specific GC B cells **(Fig. 5B, D, Fig. 7J-L)**. We posit that studying antigen valency in this way is more reflective of how B cells actively engage with real world antigens, as human B cells do not compete in a vacuum. This dramatic impact of antigen valency on B cell responses in a competitive fashion may even be reflective of how B cells naturally coevolved alongside viruses and bacteria for over 500 million years since diverging from lampreys.^82^ This may reveal insights into a possible immune evasion tactic and why some viruses avoid highly valent protein expression on their surface. HIV virions contain only 10-20 copies of Env protein on their surface^83^ and coronaviruses space their spike proteins far apart (∼25nm), tending to avoid highly repetitive organized structures.^84^

Regardless, this study highlights the importance of furthering our fundamental understanding of how B cells directly engage antigen on their surface. Greater understanding of foundational aspects of successful vaccines have furthered our abilities to design more potent and durable vaccines. One example is to simply extend antigen dosing kinetics to mimic natural antigen delivery during viral infections.^85–87^ This is in part mechanistically thought to improve antigen shuttling to GCs whilst preserving antigen structural integrity.^88^ Another is having particles that mimic viral particle sizes that can lead to extended draining kinetics, thus promoting B cell responses.^5^ Antigen multivalency, which in the context of our mosaic particles is analogous to epitope density, may have defined distances in which B cells are preferentially triggered, similar to the recently measured 20-30nm intra-fab distance.^89^ This phenomenon could be influenced by two independent surface-bound BCRs simultaneously binding one antigen or whether one BCR binding the same antigen with both Fabs. More fundamental work is needed to determine how exactly BCRs are structurally triggered on the surface of B cells post cognate antigen interaction. It is still debated whether BCRs are pulled together (e.g. crosslinked) or pulled apart to engage cell signaling following antigen binding.^90^

Our observations that antigen multivalency promoted more robust B cell responses post vaccination generally agree with previous studies.^3,5,6^ It was somewhat surprising that our analysis of epitope-specific polyclonal B cell response to our mosaic immunogens revealed little difference in the relative affinity of polyclonal B cells recruited to epitope-binding GCs on day 8 post immunization. There was only a small tendency for low valency immunogens to enrich for higher affinity B cells when compared to highest valency immunogens (12µM K_D_ vs 7.2µM K_D_ median) **(Fig. 5O, Table S5).** While this is in partial contrast with previous studies using the VRC01^gHL^ system,^6^ one explanation is that the multivalences tested in both studies were different. We could not recover a reasonable number of epitope-specific cells from mice primed with avidities lower than eOD-GT5 60mer_26_. It is plausible that we did not see a dramatic response in recruitment affinities as our “low” multivalency eOD-GT5 60mer_26_ still contains approximately 16-copies of WT eOD-GT5. Another is that we did not investigate the antigen non-binding component of the GC. This compartment is plausibly enriched for a subset of specific B cells with affinities below our detection limit of 100µM *K*_D_. However, our observed affinities were reasonably low, with several clones reporting affinities up to near the detection limit of 100µM *K*_D_, and many clones were detected at or below this limit. Our ability to identify such low affinity clones is likely due to the valency advantage that fluorescently labeled tetramers have when binding surface BCRs on B cells for cell sorting compared to monomeric measurements of *K*_D_ in SPR. In addition to presumably ultra-low affinity B cells, the antigen non-binding GC compartment contains B cells specifically responding to “dark antigen”.^79^ Dark antigen is the phrase coined for denatured or otherwise modified neo-epitopes of the immunizing antigen, with one famous example being the internal V3 loop of HIV Env.^91^ Creative future experiments will have to be designed to reveal the true nature of the antigen non-binding component of GC reactions so that these two subsets may be distinguished from one another. The fact that antigen valency promoted early affinity maturation kinetics and altered immunodominance patterns of clonal families between days 8 and 21 of the GC reaction is of interest. One possible interpretation of this is that high avidity immunogens may promote more total B cell diversity with increased absolute numbers of epitope specific B cells that are able to compete within GCs. This in turn may engender productive responses post vaccination to rare or challenging mutational pathways such as those present in some bnAb lineages.

Mosaic nanoparticles, comprising neutralizing targets for different strains of viruses such as SARS-CoV-2^24–26^ and influenza^17^ have recently been explored as a tool to preferentially prime broadly neutralizing B cell precursors following vaccination. The notion is to utilize antigen valency as a rheostat to elicit cross-protective immune responses whereby only B cells that are able to bind multiple different viral variants arrayed on the same nanoparticle are preferentially expanded in the immune response. Our study agrees with this notion and provides mechanistic insights into how such preferential expansion of desired B cells occurs *in vivo*.

Our finding that higher valency particles with fixed diameter promoted the development and maintenance of LLPCs is consistent with prior observations that this population can start to form early in an immune response and is tied to GC reactions.^92^ The imprinting hypothesis simply states that the terminal fate of a B cell, such as differentiation into a LLPC, is related to the quality of the earliest interactions with cognate antigen.^13,93^ Our data is in general agreement with this notion. However, the rescue of the VRC01^gHL^ B cell responses to low valency immunogens when placed in a mouse with restricted interclonal competition **(Fig. 6A)** potentially sheds new light on this hypothesis.^13^ One interpretation is that the “imprint” B cells experience may not be only related to the initial BCR interaction with antigen, but occur over several days, and is partially reliant on the level of early CD4 T cell help (i.e. early developing Tfh cells). Early T cell help is known to be a critical regulator of B cell entry to GCs.^35,36^ One key earlier finding was that high valency particles drive extended B:T cell contacts in the first few days post immunization.^6^ Moreover, induction of early GC or plasma cell fates by high valency particles is attenuated upon CD4 T cell depletion.^6^ Since in MD4 mice interclonal competition is substantially attenuated, the relative CD4 T cell help experienced by VRC01^gHL^ B cells may be effectively increased, contributing to the rescue of VRC01^gHL^ B cells in mice primed with low valency antigen. Moreover, a secondary interpretation is that the valencies of eOD-GT5 60mer_26_ (akin to 16-copies of WT epitope) and higher are suitable to drive VRC01-class B cell responses under attenuated interclonal competition conditions.

In sum, the results indicate that the nature of the structure of the antigen and how it interacts with BCRs on the surface of B cells is critical in driving durable and diverse immune responses post vaccination. Our study allowed us to precisely vary two independent key components of antigen avidity – affinity and multivalency. These variables, particularly multivalency, should be carefully considered when designing new vaccines.

## Limitations of the study

One limitation of the study was that the physiological relevant precursor frequencies in our model systems did limit our ability to thoroughly investigate the role of antigen valency on VRC01^gHL^ BCR diversity in both GCs and MBCs, as only the medium to high valency groups had reliably high enough cell counts for sequencing analysis. Another limitation was the inability to differentiate serum antibody responses originating from our VRC01^gHL^ or HuGL B cells compared to the endogenous response. Finally, another limitation was that these studies were done in mice. However, we used multiple preclinical mouse models that have been shown to be predictive of human B cell responses. The VRC01^gHL^-GT5 model^8^ was remarkably predictive of the human VRC01-class B cell response in the G001 clinical trial of eOD-GT8.^11^ Under high physiological affinity conditions, our model predicted that there would be an approximate 29-fold expansion from 1 in 10^6^ VRC01-class precursor frequency to B cell memory after one immunization.^8^ In the clinical trial of eOD-GT8 in humans, the observed expansion of VRC01-class precursors with a similar starting *K*_D_ and precursor frequency expanded 37-fold following one immunization.^11^ Nevertheless, it remains to be seen if our observations regarding the impacts of antigen valency on driving diverse and durable vaccine immune responses will translate to humans.

## Materials and Methods

See supplemental information for detailed methods. All animal studies were done in accordance with approved IACUC protocols.

## Author contributions

Conceptualization, W.R.S. and R.K.A.; Methodology, N.G.W., C.A.C., W.R.S., and R.K.A.; Software, N.G.W., M.V.P., L.P., and C.A.C.; Validation, N.G.W., L.P., C.A.C., and R.K.A.; Formal Analysis, N.G.W., M.V.P., L.P., C.A.C., K.G. and R.K.A.; Investigation, N.G.W., M.V.P., L.P., C.A.C., K.G., O.K., M.K., N.A., N.P., M.W., and R.K.A.; Resources, M.W., W.R.S., and R.K.A; Data Curation, N.G.W., M.V.P., L.P., C.A.C., M.W., K.G., O.K., M.K., N.A., N.P., and R.K.A.; Writing – Original Draft, R.K.A.; Writing – Reviewing & Editing, N.G.W., M.V.P., L.P., M.W., W.R.S., and R.K.A.; Visualization, N.G.W., M.W., and M.V.P.; Project Administration, N.G.W., C.A.C., W.R.S., and R.K.A.; Funding Acquisition, W.R.S. and R.K.A. Supervision, W.R.S. and R.K.A.

## Acknowledgements

R.K.A. was supported by grants 5DP2AI154410 and 5R00AI14576 as well as UTMB startup funds. N.G.W. and L.P. were supported by McLaughlin Fellowships. W.R.S. was supported by the Bill and Melinda Gates Foundation Collaboration for AIDS Vaccine Discovery (NAC INV-007522 and INV-008813), the IAVI Neutralizing Antibody Center (NAC), and National Institute of Allergy and Infectious Diseases (NIAID) UM1 Al100663 (Scripps Center for HIV/AIDS Vaccine Immunology and Immunogen Discovery) and UM1 AI144462 (Scripps Consortium for HIV/AIDS Vaccine Development). We thank Dr. Garnett Kelsoe for kindly providing the NB21-2D9 cell line. We thank Dr. Zbigniew Mikulski for helpful discussions surrounding our microscopy work. We thank Dr. Jeong Hyun Lee for helpful technical discussions regarding 10x genomics experiments and mosaic particle validation. We thank Dr. Sydney Ramirez for helpful discussions regarding 10x genomics analysis pipeline. We thank Dr. Simon Bélanger, Dr. Hugues Fausther-Bovendo, Kristy Waldrep, and Maisha Aniqua for helpful comments on the manuscript. Cell sorting and flow cytometry analysis for this project was done on instruments in the UTMB Flow Cytometry and Cell Sorting Core Lab.

## Competing Interests

W.R.S. is an inventor on patents filed by Scripps and IAVI on the eOD-GT8 monomer and 60mer immunogens. W.R.S. is an employee and shareholder of Moderna, Inc.

## Notes

### Summary of Updates

Updated figure 6 errors in labels for flow plot headers.

