## Supplementary Information for "Antigen avidity drives the durability of the vaccine immune response"

**This file includes:**

| | |
|---|---|
| Key Resources Table | Page 2 |
| Methods | Page 6 |
| Supplemental References | Page 13 |
| Figure S1: Characterization of eOD-GT5 mosaic nanoparticles and extended total GC data. | Page 14 |
| Figure S2: Extended VRC01$^{gHL}$ mutation data. | Page 16 |
| Figure S3: Extended VRC01$^{gHL}$ MBC data. | Page 18 |
| Figure S4: Extended VRC01$^{gHL}$ MBC data from day 42 post-boost. | Page 20 |
| Figure S5: Extended endogenous response data. | Page 22 |
| Figure S6: Adjuvant, dosing, and extended MD4 model data. | Page 24 |
| Figure S7: Characterization of eOD-GT8 mosaic nanoparticles and extended HuGL18 and HuGL17 data. | Page 26 |
| Table S1: Co-transfection ratios and KinExa data for eOD-GT5 mosaic nanoparticles. | Page 28 |
| Table S2: Number of amino acid mutations in VRC01$^{gHL}$ GC and MBC sequences. | Page 29 |
| Table S3: Simpon's Index of Diversity for VRC01$^{gHL}$ GC and MBC BCR sequencing. | Page 30 |
| Table S4: Simpon's Index of Diversity for Endogenous CD4bs WT GC B cells. | Page 31 |
| Table S5: Quantification of affinity from antibodies derived from cultured GC B cells. | Page 32 |
| Table S6: Co-transfection ratios and KinExa data for eOD-GT8 mosaic nanoparticles. | Page 33 |

**Key resources table**

| REAGENT or RESOURCE | SOURCE | IDENTIFIER |
|---|---|---|
| Antibodies | | |
| Rat monoclonal anti-mouse-FoxP3 Alexa Fluor 647 (clone MF-14) | Biolegend | Cat #:126408 |
| Mouse monoclonal anti-mouse-CD45.1 Alexa Fluor 647, Brilliant Violet 711, or FITC (clone A20) | Biolegend | Cat #:110720, 110739, 110706 |
| Armenian hamster anti-mouse-TCRβ chain Alexa Fluor 488 (clone H57-597) | Biolegend | Cat #:109215 |
| Rat monoclonal anti-mouse-CD38 APC/Cyanine7 or PE/Cyanine7 (clone 90) | Biolegend | Cat #:102727, 102718 |
| Rat monoclonal anti-mouse-CD4 APC/Fire750, FITC, or BUV395 (clone GK1.5) | Biolegend / BD Biosciences | Cat #:100460, 100406 / 563790 |
| Rat monoclonal anti-mouse-CD8a APC/Fire750 (clone 53-6.7) | Biolegend | Cat #:100766 |
| Rat monoclonal anti-mouse-CD185 (CXCR5) Brilliant Violet 421 (clone L138D7) | Biolegend | Cat #:145511 |
| Armenian hamster monoclonal anti-mouse-CD80 Brilliant Violet 421 (clone 16-10A1) | Biolegend | Cat #:104726 |
| Rat monoclonal anti-mouse/human-CD45R/B220 Brilliant Violet 421 or Brilliant Violet 785 (clone RA3-6B2) | Biolegend | Cat #:103240, 103246 |
| Rat monoclonal anti-mouse-CD90.2 (Thy1.2) Brilliant Violet 510 (clone 53-2.1) | Biolegend | Cat #:140319 |
| Rat monoclonal anti-mouse-IgD Brilliant Violet 510, Brilliant Violet 650, or FITC (clone 11-26c.2a) | Biolegend | Cat #:405723, 405721, 405704 |
| Rat monoclonal anti-mouse-CD279 (PD-1) Brilliant Violet 605 (clone 29F.1A12) | Biolegend | Cat #:135219 |
| Rat monoclonal anti-mouse-CD73 Brilliant Violet 605 (clone TY/11.8) | Biolegend | Cat #:127215 |
| Rat monoclonal anti-mouse-CD138 (Syndecan-1) Brilliant Violet 650 (clone 281-2) | Biolegend | Cat #:142518 |
| Mouse monoclonal anti-mouse-CD45.2 Brilliant Violet 711 or FITC (clone 104) | Biolegend | Cat#:109847, 109806 |
| Rat monoclonal anti-mouse/human-CD44 Brilliant Violet 711 (clone IM7) | Biolegend | Cat #:103057 |
| Rat monoclonal anti-mouse-CD273 (B7-DC, PD-L2) PE or PE/Cyanine7 (clone TY25) | Biolegend | Cat #:107206, 107214 |
| Syrian hamster monoclonal anti-mouse-CD278 (ICOS) PE (clone 15F9) | Biolegend | Cat #:107705 |
| Rat monoclonal anti-mouse-GL7 (T and B cell Activation Marker) PE, PerCP/Cyanine5.5, or BB700 (clone GL7) | Biolegend | Cat #:144608, 144610, 624409 |
| Zombie NIR Fixable Viability Kit | Biolegend | Cat #:423106 |
| Zombie Yellow Fixable Viability Kit | Biolegend | Cat #:423103 |
| Rat monoclonal anti-mouse-CD43 FITC (clone S7) | BD Biosciences | Cat #:553270 |
| Rat monoclonal anti-mouse-IgG1 PE-CF594 (clone A85-1) | BD Biosciences | Cat #:562559 |
| Rat monoclonal anti-mouse-IgM BUV737 (clone II/41) | BD Biosciences | Cat #:749308 |
| TotalSeq™-C0301 anti-mouse Hashtag 1 | Biolegend | Cat #:155861 |
| TotalSeq™-C0302 anti-mouse Hashtag 2 | Biolegend | Cat #:155863 |
| TotalSeq™-C0303 anti-mouse Hashtag 3 | Biolegend | Cat #:155865 |
| TotalSeq™-C0304 anti-mouse Hashtag 4 | Biolegend | Cat #:155867 |
| TotalSeq™-C0305 anti-mouse Hashtag 5 | Biolegend | Cat #:155869 |

| | | |
|---|---|---|
| Goat anti-mouse IgG (Fc)-HRP | Bethyl Labs | Cat #:A90-131P |
| Goat anti-mouse Ig - UNLB | Southern Biotech | Cat #:1010-01 |
| Chemicals, peptides, and recombinant proteins | | |
| eOD-GT5 WT 60mer | Produced in house | N/A |
| eOD-GT5 mosaic WT-KO 10-1 60mer | Produced in house | N/A |
| eOD-GT5 mosaic WT-KO 1-1 60mer | Produced in house | N/A |
| eOD-GT5 mosaic WT-KO 1-5 60mer | Produced in house | N/A |
| eOD-GT5 mosaic WT-KO 1-10 60mer | Produced in house | N/A |
| eOD-GT5 KO 60mer | Produced in house | N/A |
| eOD-GT8 WT 60mer | Produced in house | N/A |
| eOD-GT8 mosaic WT-KO 10-1 60mer | Produced in house | N/A |
| eOD-GT8 mosaic WT-KO 1-1 60mer | Produced in house | N/A |
| eOD-GT8 mosaic WT-KO 1-5 60mer | Produced in house | N/A |
| eOD-GT8 mosaic WT-KO 1-10 60mer | Produced in house | N/A |
| eOD-GT8 KO 60mer | Produced in house | N/A |
| eOD-GT5 avi tagged monomer | Produced in house | N/A |
| eOD-GT5 KO avi tagged monomer | Produced in house | N/A |
| eOD-GT8 avi tagged monomer | Produced in house | N/A |
| eOD-GT8 KO avi tagged monomer | Produced in house | N/A |
| Streptavidin-APC | Biolegend | Cat #:405243 |
| Streptavidin-PE | Biolegend | Cat #:405204 |
| Recombinant mouse IL-4 | Peprotech | Cat #:214-14 |
| Critical commercial assays | | |
| Fetal bovine serum, value | Gibco | Cat #:A52567-01 |
| Fetal bovine serum, qualified | Gibco | Cat #:26140-079 |
| Horse serum | HyClone | Cat #:SH30074.03 |
| Dulbecco's PBS without calcium and magnesium | Gibco | Cat #:14190-250 |
| QuadroMACS separator | Miltenyi Biotec | Cat #:130-091-051 |
| LS columns | Miltenyi Biotec | Cat #:130-042-401 |
| Anti FITC microbeads | Miltenyi Biotec | Cat #:130-048-701 |
| Histopaque-1077 | Sigma-Aldrich | Cat #:10771-100ML |
| Pen Strep | Gibco | Cat #:15140-122 |
| 2-Mercaptoethanol | Gibco | Cat #:21-985-023 |
| HEPES | Corning | Cat #:25-060-CI |
| Sodium pyruvate | Sigma-Aldrich | Cat #:S8636-100ML |
| MEM Nonessential Amino Acids 100X | Corning | Cat #:25-025-CI |
| 0.5%, 10X Trypsin-EDTA | Invitrogen | Cat #:15-4000-54 |
| PBS, 1X, without calcium and magnesium | Corning | Cat #:21-040-CV |
| Sodium azide | Sigma-Aldrich | Cat #:S2002-100G |
| RPMI-1640 with L-Glutamine | Gibco | Cat #:11875-093 |
| Dulbecco's Modified Eagle Medium with 4.5 g/L D-glucose and L-Glutamine | Gibco | Cat #:11965-092 |
| Fluoromount-G Slide Mounting Medium | Southern Biotech | Cat #:0100-01 |
| BSA | Fisher Scientific | Cat #: BP1600-1 |
| Tween 20 | Sigma-Aldrich | Cat #:P7949-500ML |
| NUNC Maxisorp plates | Thermo Scientific | Cat #:442404 |
| TMB substrate reagent kit | BD Biosciences | Cat #:555214 |
| Hard-Shell High Profile 96-Well Semi-Skirted PCR Plates | Bio-Rad | Cat #:HSS9601 |
| RNase Zap | Invitrogen | Cat #:AM9782 |
| RNase inhibitor, murine | New England Biolabs | Cat #:M0314L |
| Poly(A), Polyadenylic acid | Millipore Sigma | Cat #:10108626001 |

| | | |
|---|---|---|
| Trizma hydrochloride solution, pH 8.0 | Sigma-Aldrich | Cat #:T2694-100ML |
| Ambion Nuclease-Free Water | Invitrogen | Cat #:AM9937 |
| ACK lysing buffer | Gibco | Cat #:A10492-01 |
| Propidium iodide | Biolegend | Cat #:421301 |
| UltraComp eBeads Plus Compensation Beads | Invitrogen | Cat #:01-3333-42 |
| ArC Negative Beads | Invitrogen | Cat #:A10346B |
| ArC Reactive Beads | Invitrogen | Cat #:A10346A |
| eBioscience FOXP3/Transcription Factor Staining Buffer Kit | Invitrogen | Cat #:00-5523-00 |
| BD Cytofix/Cytoperm Fixation/Permeabilization Kit | BD Biosciences | Cat #:554714 |
| Alhydrogel | InvivoGen | Cat #:vac-alu-250 |
| AddaS03-like adjuvant | InvivoGen | Cat #:vac-as0310 |
| Sigma (RIBI) Adjuvant System | Millipore Sigma | Cat #:S6322-1VL |
| Chromium Next GEM Single Cell 5’ Kit v2, 16 rxns | 10X Genomics | Cat #:1000263 |
| Chromium Next GEM Chip K Single Cell Kit, 48 rxns | 10X Genomics | Cat #:1000286 |
| Dual Index Kit TT Set A, 96 rxns | 10X Genomics | Cat #:1000215 |
| Dual Index Kit TS Set A, 96 rxns | 10X Genomics | Cat #:1000251 |
| Chromium Single Cell Mouse BCR Amplification Kit, 16 rxns | 10X Genomics | Cat #:1000255 |
| Dual Index Kit TN Set A, 96 rxns | 10X Genomics | Cat #:1000250 |
| Chromium GEM-X Single Cell 5’ Kit v3, 16 rxns | 10X Genomics | Cat #:1000699 |
| Chromium GEM-X Single Cell 5’ Chip Kit v3, 4 chips | 10X Genomics | Cat #:1000698 |
| 5’ Feature Barcode Kit, 16 rxns | 10X Genomics | Cat #:1000541 |
| Deposited data | | |
| Experimental models: Cell lines | | |
| NB-21.2D9 cell line | Kuraoka et al., 2016[1] | N/A |
| Experimental models: Organisms/strains | | |
| Mouse: B6.SJL-*Ptprc*$^a$*pepc*$^b$/BoyJ | The Jackson Laboratory | JAX: 002014 |
| Mouse: C57BL/6J | The Jackson Laboratory | JAX: 000664 |
| Mouse: C57BL/6-Tg(IghelMD$)4Ccg/J | The Jackson Laboratory | JAX: 002595 |
| Mouse: B6.Cg-Tg(Prdm1-EYFP)1Mnz/J | The Jackson Laboratory | JAX: 008828 |
| Mouse: VRC01$^{gH+/+gL+/+}$ | Abbott et al., 2018[2] | N/A |
| Mouse: VRC01$^{gH+/-L+/-}$ | Abbott et al., 2018[2] | N/A |
| Mouse: HuGL18 | Huang et al., 2020[3] | N/A |
| Mouse: HuGL17 | Huang et al., 2020[3] | N/A |
| Oligonucleotides | | |
| Primer 1mFH_VII: CCTGTCAGTAACTRCAGGTGTCC | IDT | Custom ordered |
| Primer 1mRG* (Gamma): AGAAGGTGTGCACACCGCTGGAC | IDT | Custom ordered |
| Primer 1mFK_I: RGTGCAGATTTTCAGCTTCCTGCT | IDT | Custom ordered |
| Primer 1mRK*: ACTGAGGCACCTCCAGATGTT | IDT | Custom ordered |
| Primer 2mFG*: GGGAATTCGAGGTGCAGCTGCAGGAGTCTGG | IDT | Custom ordered |

| | | |
|---|---|---|
| Primer 2mRG* (Gamma): GCTCAGGGAARTAGCCCTTGAC | IDT | Custom ordered |
| Primer 2mFK*: GAYATTGTGMTSACMCARWCTMCA | IDT | Custom ordered |
| Primer 2mRK*: TGGGAAGATGGATACAGTT | IDT | Custom ordered |
| Software and algorithms | | |
| Flowjo v10 | Treestar | https://www.flowjo.com/ |
| Prism X | GraphPad | https://www.graphpad.com/ |
| Microsoft Office | Microsoft | https://www.office.com/ |
| Adobe Creative Cloud VL 6 | Adobe | https://www.adobe.com/creativecloud.html |
| QuPath-0.5.0 | QuPath | https://qupath.github.io/ |
| Geneious Prime 2025.0 | Dotmatics | https://www.geneious.com/ |
| Cell Ranger v9 | 10X Genomics | https://www.10xgenomics.com/support/software/cell-ranger/latest |
| Sequencing Analysis and Data Library for Immunoinformatics Exploration (SADIE) | Github | https://github.com/jwillis0720/sadie |
| RStudio-2024.12.1-563 | Posit Software | https://posit.co/download/rstudio-desktop/ |
| Other | | |

#### METHODS

##### Protein Production

His-Avi-tagged monomeric eOD-GT5, eOD-GT5 KO11b, eOD-GT8, and eOD-GT8 KO11 were produced, purified, and biotinylated as previously described[4] for use in flow cytometry and ELISA experiments. Non-biotinylated monomeric eOD-GT5 and eOD-GT8 were similarly produced and purified for KinExa and SPR experiments. Nanoparticle 60mer immunogens were generated via transient transfection of HEK-293F cells (ThermoFisher) using 1 mg of total DNA per liter of cultured cells and 3 mg of polyethylenimine transfection reagent. Mosaic nanoparticles were produced by co-transfecting cells with varying ratios of plasmids encoding WT and KO nanoparticles.

Immunogens were purified using Galanthus nivalis lectin affinity chromatography (Vectorlabs), followed by size exclusion chromatography with a Superose 6 16/600 PG column (Cytiva). The molecular weight and homogeneity of the antigens were assessed using size-exclusion chromatography–multi-angle light scattering (SEC-MALS) in phosphate-buffered saline (PBS). Analyses were conducted with Superdex 75 10/300 GL columns for monomers and Superose 6 Increase 10/300 GL columns for nanoparticles (Cytiva), operating at an isocratic flow rate of 0.5 mL/min. Detection was performed using DAWN HELEOS II and Optilab T-rEX detectors (Wyatt Technology). Endotoxin levels in immunogen preparations were confirmed to be <5 EU/mg using an Endosafe instrument (Charles River).

Nanoparticle immunogens assessed for quality using transmission electron microscopy. Briefly, samples were deposited onto glow-discharged, carbon-coated copper mesh grids, stained with uranyl formate, and imaged using a Talos L120C (ThermoFisher) electron microscope at a nominal magnification of 120,000× with a CETA 16M CMOS camera (ThermoFisher).

Human IgG1 and Fab versions of VRC01 and GL-VRC01 were produced and purified as previously described.[5] Murine IgG2a versions of VRC01 and GL-VRC01 were kindly provided by Sonya Haupt (La Jolla Institute for Immunology). Naked lumazine synthase was produced by BlueSky BioServices as previously described.[3]

##### Kinetic Exclusion Assay (KinExA)

Mature VRC01 Fab and GL-VRC01 Fab were used as Constant Binding Partners (CBP), for the eOD-GT5 and eOD-GT8 experiments respectively. Monomeric eOD-GT5 and eOD-GT8 were dialyzed overnight against 1x HBS-EP+ buffer (Teknova H8022) without BSA and coupled to UltraLink azlactone-activated, beaded-polyacrylamide resin (ThermoFisher 53110). The cross-linking reaction was carried out in 0.5 M Sodium Citrate buffer pH 6.5 with overnight incubation. Running buffer was 1x HBS-EP+ pH 7.4 supplemented with BSA (Sigma A3294) at 1mg/mL. The labeling antibody was Alexa Fluor 647 conjugated AffiniPure Fab Fragment Goat Anti-

Human IgG (H+L), (Jackson ImmunoResearch 109-607-003), used at 1000 ng/mL in running buffer (prepared fresh from 1000x concentrated stock that is stored at -80C). Experiments were run on a KinExA 3200 with Autosampler model SAPIDYNE-AIM3300 (Sapidyne Instruments). We ran all measurements and equilibrations at room temperature (about +25 C). The nanoparticle antigens were serially diluted into samples having constant concentration of Fab. All samples were equilibrated for 24 hours before the start of the measurement. All data points were measured in duplicates. We analyzed the data using n-curve function of the KinExA Pro software (Version 4.0.12) to determine $K_D$ of the nanoparticle antigens for the CBP Fab and Titrant % Activity for each nanoparticle antigen. Normalized % Activity was calculated by dividing the Titrant % Activity for each mosaic nanoparticle antigen by the Titrant % Activity for the WT nanoparticle antigen.

**ELISA for validation of particle mosaicism**

Costar Maxisorp plates were coated with 2μg/ml human germline VRC01 mAb at RT in 1XPBS. Plates were then blocked overnight at 4°C (0.5% BSA/PBS) and 0.5μg/mL of eOD-GT 60mers in blocking buffer were added one hour. Previously screened KO+WT- nojima culture supernatants (described below) diluted 1:1, were added for one hour. Anti-mouse IgG-HRP was utilized for detection (Bethyl Labs). Plates were washed between all steps and subsequently visualized using TMB substrate (Thermo Scientific), stopped using 0.2M $H_2SO_4$, and read on a BioTek Synergy H1 plate reader. LOD was determined by blank wells lacking either nanoparticles or both nanoparticles and KO+WT- serum.

**Mice and Immunizations**

B6.SJL-*Ptprc*$^{a}$*pepc*$^{b}$/BoyJ mice (CD45.1$^{+/+}$) were purchased from the Jackson labs (strain #002014) (Bar Harbor, ME) and maintained as an internal breeding colony at the University of Texas Medical Branch (UTMB) to be used as recipient mice. C57BL/6J mice were also purchased from the Jackson labs (strain #000664) to be used as recipient mice for experiments. Both males and females between 8-16 weeks of age were used for experiments. For any given individual experiment, age and sex were specifically matched. VRC01$^{gHL}$ mice were maintained from original colony developed at the La Jolla Institute (LJI) as previously described[2] at UTMB. HuGL18 and HuGL17 mice were maintained from original colony developed at the Scripps Research Institute (TSRI) as previously described[2,3] at UTMB. C57BL/6-Tg(IghelMD4)4Ccg/J mice (MD4) were purchased from the Jackson labs (strain #002595) and maintained as an internal breeding colony at UTMB, bred to C57BL/6J mice. B6.Cg-Tg(Prdm1-EYFP)1Mnz/J mice (Blimp1 YFP) were purchased from the Jackson labs (strain #008828) and maintained as an internal breeding colony at UTMB, backcrossed 1 generation to C57BL/6J mice and then bred to VRC01$^{gHL}$ mice. Preparations of given immunogens (eOD-GT5 or -8 60mers and respective mosaic nanoparticles) were diluted in PBS (200 μg/ml for 100 μl/20 μg/mouse and mixed at a 1:1 ratio with 100 μl/mouse Alhydrogel 2% (Invivogen)) for at least 20 minutes and then injected intraperitoneally (i.p.) (total volume 200 µl). At day of spleen or bone marrow collection, the peritoneal cavity of all immunized mice was inspected for presence of Alum, assuring proper injection of mice. All mice were housed under specific pathogen free conditions at UTMB. All mouse experiments were done under approved Institutional Animal Care and Use Committee protocols at UTMB.

**Adoptive B Cell Transfer**

Lymphocytes were prepared from spleens of heterozygous VRC01$^{gHL}$, HuGL18, HuGL17, or Blimp1$^{YFP+}$ VRC01$^{gHL}$ mice by density gradient centrifugation (Histopaq, Sigma). B cells were purified by negative selection by magnetic depletion using FITC conjugated anti-CD43 (clone S7), anti-FITC microbeads (Miltenyi Biotec), and an LS column (Miltenyi Biotec). LS columns were washed once with 8 ml of buffer which was discarded prior to running cells. Collected cells were then checked by flow cytometry for B cell purity and antigen-positive cells using fluorescently labeled eOD-GT5 (VRC01$^{gHL}$) or eOD-GT8 (HuGL) prior to transfer. Cells were enumerated on a hemocytometer and transferred retro-orbitally (RO) into recipient hosts. All steps were performed in transfer buffer (5% horse serum/Dulbecco's PBS (with calcium and magnesium)). All tubes, pipette tips, and syringe needles were pre-coated in transfer buffer for at least one hour at room temperature in transfer buffer to minimize cell loss. All B cell transfers were done by either NGW or RKA, and extensive validation of B cell take and actual precursor frequency in recipients was done to ensure consistency prior to this study. Mice were rested for approximately 24 hours before immunization.

**Flow Cytometry**

Single cell suspensions were generated by mechanical dissociation of spleens and bone marrow. Red blood cells were removed through ACK lysis. Cells were prepared in FACS buffer (5% FCS/PBS), enumerated, and Fc blocked (clone 2.4G2, BD Biosciences). Mixtures of mAbs from Biolegend and BD Biosciences were added for 30 minutes at 4°C. Cells were then washed and fixed in either Foxp3 fixation kit (eBioscience) or cytofix/cytoperm (BD Biosciences). If appropriate, secondary intracellular stains were performed the next day. Samples were acquired on a NovoCyte Penteon (Agilent). Samples were analyzed on FlowJo v10.10.0. For most experiments, 1/3 of the spleen was snap frozen for histological analysis and the remaining 2/3 was counted and processed for flow cytometry analysis. Absolute numbers of cells per spleen were calculated by hemocytometer counts from experiments utilizing whole spleens for flow cytometry. For memory B cell and long-lived plasma cell timepoints 10 million total events, or about 3-4 million B cells, were collected. Rare VRC01$^{gHL}$ populations were confirmed by comparison to negative naïve controls, as well as confirmation by back gating. Limit of detection (LOD) was taken as the average of all naïve control frequencies for the specific gated populations.

**Single cell sorting for VRC01 B cell analysis**

Single cell suspensions were generated from spleens and bone marrow by mechanical dissociation and ACK lysis was used to remove red blood cells. Cells were prepared in FACS buffer (5% FCS/PBS) and Fc blocked (clone 2.4G2, BD Biosciences). mAb cocktails from Biolegend and BD Biosciences were added for 30 minutes at 4°C. Samples were acquired on a FACSAria Fusion (BD Biosciences) with four-way purity setting enabled at pressure 70 psi using a 70µM nozzle and sorted into wells of 96-well plates containing 15 µl of lysis buffer cocktail (ambion nuclease-free water (Life Technologies), trizma (0.1M), poly A (10 µg/mL), and RNase inhibitor (500 U/mL)). Cells were spun down, frozen on dry ice, and stored at -80°C until used for sequencing. First strand cDNA synthesis was carried out using SuperScript II Reverse Transcriptase (Invitrogen) according to instruction from manufacturer. Nested PCR was performed in a total volume of 16 uL with HotStar Taq Master Mix (Qiagen) using IgG- and IgK- specific primer pools and thermocycling conditions described previously.[6] Secondary nested PCR reactions were carried out using Phusion polymerase (ThermoFisher) with forward primers specific for the VRC01 HC and LC leader sequences as described previously.[2] PCR products were run on 1 or 1.5% agarose gel to confirm proper amplification. Wells with amplicons of correct size were sent to Eton Biosciences for Sanger sequencing. Sequencing primer selection was as described previously.[2]

**Single cell sorting for endogenous B cell analysis**

Single cell suspensions were generated following the method above. TotalSeq hashtag antibodies were added to each individual cell suspension prior to sorting for sample deconvolution of resultant sequences. Samples were acquired on a FACSAria Fusion (BD Biosciences) with single cell setting enabled at pressure 45 psi using a 85µM nozzle. Cells of each group were sorted into 10uL of pure FCS (Gibco) up to 25,000 cells per group. Cell suspensions were then spun down and maintained on ice until utilized in the 10x Genomics v2 or v3 GEM reaction.

**NGS sequencing of endogenous BCR libraries**

Libraries containing indexed V(D)J and Feature barcode sequences were generated according to the Chromium Next GEM Single Cell 5’ v2 or v3(Dual Index) User Guide. Libraries were pooled and sequenced on a NextSeq2000 (Illumina) sequencer.

**BCR analysis pipeline**

VRC01$^{gHL}$ BCR sequences were quality-checked and trimmed in Genius Prime 2024.0.7. Heavy chain and light chain sequences were aligned using the reference amino acid sequences of unmutated VRC01$^{gH}$ and VRC01$^{gL}$, respectively. Mutation frequencies were calculated using sequences that were trimmed and aligned as the denominator. VRC01-class key mutations from paired sequences in individual mice were scored as described previously and their median counts taken.[7]

Data from NGS sequencing was processed using 10x Genomics CellRanger v9 VDJ and Count functions to create FastA files. FastA files from both NGS and paired sanger sequences were analyzed using the **S**equencing **A**nalysis and **D**ata Library for **I**mmunoinformatics **E**xploration (SADIE). SADIE utilizes IgBLAST for the analysis of nucleotide sequences to name and call mutations against endogenous germline encoded murine Ig genes and the human Ig genes present in the VRC01$^{gHL}$ mouse. BCR clustering was performed using the SADIE cluster module with average-linkage clustering. Individual sequences within a cluster were required to come from the same animal and to share the same heavy and light V gene calls and the same CDRH3 lengths. The distance between two antibodies was the sum of Levenshtein distances between all of the CDRs. An Average-linkage clustering with a distance cutoff of 5 was used for VRC01$^{gHL}$ paired sequences, whereas a distance cutoff of 2 was used for endogenous sequences. VRC01$^{gHL}$ sequences were grouped by timepoint and nanoparticle prior to clustering in Fig. 2H-J and grouped by individual mice in Fig. 2K-L.

Endogenous total projected unique clusters in Fig. S5Q were calculated by taking the frequency of clonal clusters of recovered paired BCR clones multiplied by the total number of sorted cells.

**Nojima Single B Cell Cultures**

Single B cells were cultured in the presence of NB-21.2D9 feeder cells (G Kelsoe). NB-21.2D9 cells were seeded into 96-well plates at 1,000 cells/well in B cell media [BCM] (RPMI-1640 [Invitrogen] supplemented with 10% Qualified FBS (Gibco), 5.5 x $10^{-5}$ M 2-mercaptoethanol, 10 mM HEPES, 1 mM sodium pyruvate, 100 units/mL penicillin, 100 µg/mL streptomycin, and MEM nonessential amino acid (Invitrogen). The next day (day 0), recombinant mouse IL-4 (Peprotech; 2 ng/mL) was added to the cultures, and then single B cells were directly sorted into each well by a FACSAria Fusion (BD Biosciences). On day 2, 50% of the culture media volume was removed and 100% volume of fresh BCM was added to the cultures. On days 3-8, two-thirds of the culture media were replaced with fresh BCM every day. On day 10, culture supernatants were harvested for ELISA determinations and culture plates were stored at -80°C for V(D)J amplifications.

**Enzyme-linked Immunosorbent Assays (ELISA)**

Costar Maxisorp plates were coated with 0.5µg/ml of either eOD-GT 60mers (either WT or KO from either GT5 or GT8), 2µg/ml lumazine synthase, or total Ig (Southern Biotech) for one hour at RT in 1X PBS. Plates were then blocked for at least 1 hour at RT or overnight at 4°C (0.5% BSA/PBS) and serially diluted serum samples, or culture supernatants diluted 1:1, were added for one hour. Anti-mouse IgG-HRP was utilized for detection (Bethyl Labs). Plates were washed between all steps and subsequently visualized using TMB substrate (Thermo Scientific), stopped using 0.2M $H_2SO_4$, and read on a BioTek Synergy H1 plate reader. Area under the curve (AUC) was calculated for each mouse. CD4bs-WT epitope specific responses, delta AUC, was calculated by subtracting the KO-specific AUC from the WT-specific AUC for each mouse.[8]

**Surface Plasmon Resonance**

Nojima culture supernatants were screened for total Ig by ELISA (described above). Wells above the LOD were selected for SPR analysis. LOD was calculated as one standard deviation above the average of blank wells.

We measured the kinetics and affinities of antibody-antigen interactions using a Carterra LSA instrument with HC30M sensor chips (Carterra) and 1× HBS-EP+ (pH 7.4) running buffer supplemented with 1 mg/mL BSA. Chip surface preparation followed Carterra software guidelines for ligand capture. Capture antibody Goat Anti-Mouse IgG Fc (SouthernBiotech 1013-01) was amine-coupled at 3000–4000 RU in 10 mM sodium acetate (pH 4.5). Regeneration was performed using 1.7% phosphoric acid with a 60-second contact time, injected three times per cycle.

Ligands were provided as B-cell/Nojima supernatants in 96-well plates (200 µL per well) and transferred to 1 mL 96-well plates containing 120 µL of running buffer per well. Ligand capture had a contact time of 5 minutes. Analyte samples were buffer exchanged via dialysis and quantified using a NanoDrop 2000c spectrophotometer at 280 nm. Raw sensograms were analyzed with Carterra Kinetics software using interspot and blank double referencing and a Langmuir model.

We covered a broad range of affinities, adjusting referencing practices based on ligand off-rates. For fast off-rates (>0.009 $s^{-1}$), automated batch referencing with overlay y-aline was used. For slow off-rates (≤0.009 $s^{-1}$), manual process referencing with serial y-aline was applied. After automated analysis, additional filtering was performed to exclude datasets where the highest response signals were lower than negative controls. This filtering was automated using an R script as previously described.[4]

Final datasets consisted of kinetic and affinity measurements of antibodies derived from Nojima culture supernatants, evaluated against both eOD-GT5 WT and KO analytes. To ensure that only mAbs binding the CD4bs $WT^{+}KO^{-}$ epitope were being analyzed, mAbs exhibiting measurable binding (<100µM $K_D$) to GT5 KO analytes, even if they bound GT5 WT, were excluded from analysis. Additionally, WT-specific responses at or below the assay limit of detection were excluded from analysis. Only interactions demonstrating specific, above-threshold binding to WT analyte with no detectable binding to KO were retained for downstream analysis.

**Histological Analysis**

Spleens were frozen in Tissue-Tek (Electron Microscopy Sciences) in a liquid nitrogen-cooled bath of 2-methylbutane. At least 50 5 µm-thick cryosections were cut from each spleen from each time point at least 100 µm apart in a cryostat set to -22°C. Sections were then adhered to RT SuperFrost Plus slides (Fisher Scientific), air-dried for 2 hours at RT, and fixed in a 1:1 mixture of acetone and methanol for 10 minutes at -30°C. Sections were covered with Epredia Shandon plastic coverplates (Fisher Scientific) and rehydrated and blocked in 0.5% BSA/0.1% Tween-20/PBS (stain/wash buffer used in all subsequent steps) in a Freequenza rack box (UTMB 3D

Printing Core). Sections were treated with Fc block at 1:50 (clone 2.4G2, BD Biosciences) and naïve Bl/6 serum at 1:5 for 30 minutes at RT and stained with mAbs at 1:200 each for 1 hour at RT [B220-(BV421) clone RA3-6B2, GL7-(PE), TCRβ-(A488) clone H57-597,CD45.1-(A647) clone A20] (Biolegend, San Diego CA). Slides were then washed 3 times, mounted in ProLong Gold™ Antifade Mountant (Invitrogen), and imaged using a Zeiss Axio Scan.Z1 slide scanner with Zen acquisition software. QuPath version 0.5.0 was used for all subsequent analysis.[9] The algorithm for GCs and CD45.1$^+$ was based on a pixel classifier and was trained on representative images with dedicated annotations for the GL7 and CD45.1 channels, respectively. In the GL7 channel, holes were filled and all GL7$^+$ areas (> 1,000 μm$^2$) were identified as potential GCs. All selected GCs were then evaluated as either appropriate or noise. In the CD45.1 channel, holes were filled, and exact areas (> 50 μm$^2$) were identified and evaluated as appropriate or noise. GCs from the day 20 timepoint of mice receiving eOD-GT5 KO were used to determine the LOD for detecting CD45.1 within GCs. The exact CD45.1$^+$ percent area, or VRC01$^{gHL}$ cell occupancy, was calculated as a fraction of each identified GC.

**Statistical Analysis and Graphs**

Unless stated otherwise, all statistical analysis were calculated in Prism v10.4.1 or R v4.3.0. The respective statistical tests used are indicated in the legends of corresponding figures. Unless otherwise stated, comparison of groups was accomplished by Kruskal-Wallis with Dunn's multiple comparison's correction with an alpha of 0.05, *$p<0.05$, **$p<0.01$, ***$p<0.001$, ****$p<0.0001$. Correlations between percent normalized activity of nanoparticles and cell frequencies were calculated by a Spearman coefficient analysis. All graphs were generated in Prism. Zero values are depicted below LOD on graphs with log scales on the y axis. Statistics were run on actual values. Bubble graphs depicting clonal families were generated in R using the packingcircles package.

Simpson's index of diversity ($D$) was calculated by the following equation:

$$D = 1 - \left(\frac{\Sigma\, n(n-1)}{N(N-1)}\right)$$

n = total number of sequences belonging to an individual cluster

N = total number of clusters

### Supplementary Figure 1

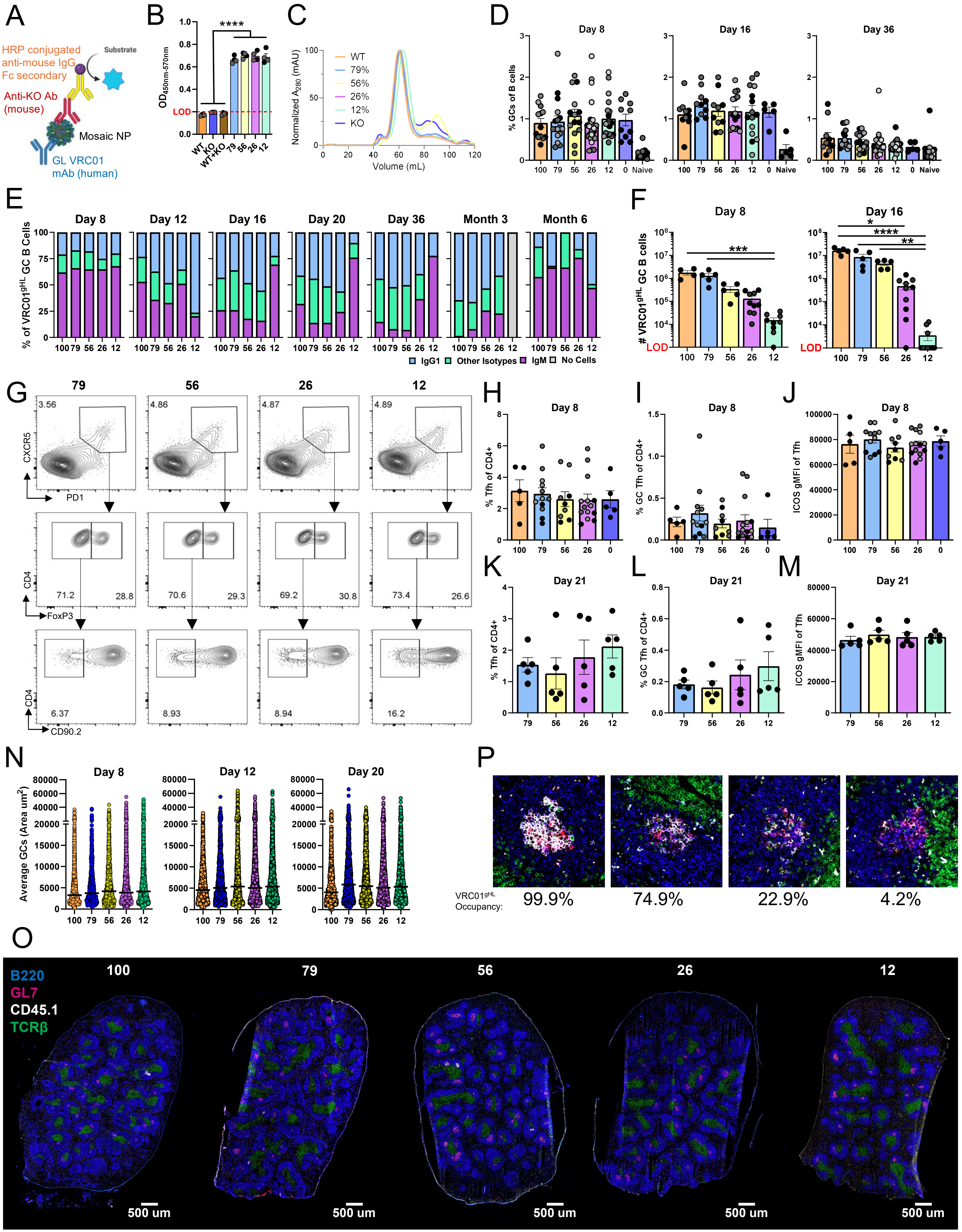

**Figure S1. Characterization of eOD-GT5 mosaic nanoparticles and extended total GC data, related to Figure 1 and Table S1. A)** Schematic of ELISA used to validate mosaic nanoparticles. **B)** Results of ELISA shown in A, demonstrating that only GT5 mosaic nanoparticles bind both CD4bs WT and KO specific antibodies, compared to fully WT or KO 60mer nanoparticles. Each dot indicates technical replicate, shades of grey indicate different preps of particles. Mann-Whitney U test comparing pooled non-mosaic particle to mosaic particle groups. ****p<0.0001. **C)** Normalized SEC profiles of eOD-GT5 mosaic nanoparticles. **D)** Quantification of total GC frequency in mice immunized with mosaic nanoparticles. **E)** Isotype distribution of responding VRC01$^{gHL}$ GC B cells. **F)** Quantification of total VRC01$^{gHL}$ GC B cell numbers in mice immunized with mosaic nanoparticles. Isolated from whole spleen. Kruskal-Wallis with Dunn's correction. *p<0.05, **p<0.01, ***p<0.001, ****p<0.0001. **G)** Representative flow cytometry from day 21 to quantify Tfh populations pre-gated on SSL/B220-/CD4+. **H-M)** Quantification of Tfh profiles on days 8 and 21. **(H,K)** Total Tfh (PD1+/CXCR5+/Foxp3-), **(I, L)** GC Tfh (CD90.2- Tfh), and **(J, M)** gMFI of Tfh ICOS expression. **N)** Quantification of average GC size in spleen by area. Bar indicates median. **O)** Representative total half spleen histological images from day 12. **P)** Representative histological images from day 12 from various groups demonstrating percent area of VRC01$^{gHL}$ cells within GCs. **(B, F)** LOD = limit of detection. **(B, D, F, H-M)** Mean ± SEM. Shades of grey dots indicate independent experiments.

### Supplementary Figure 2

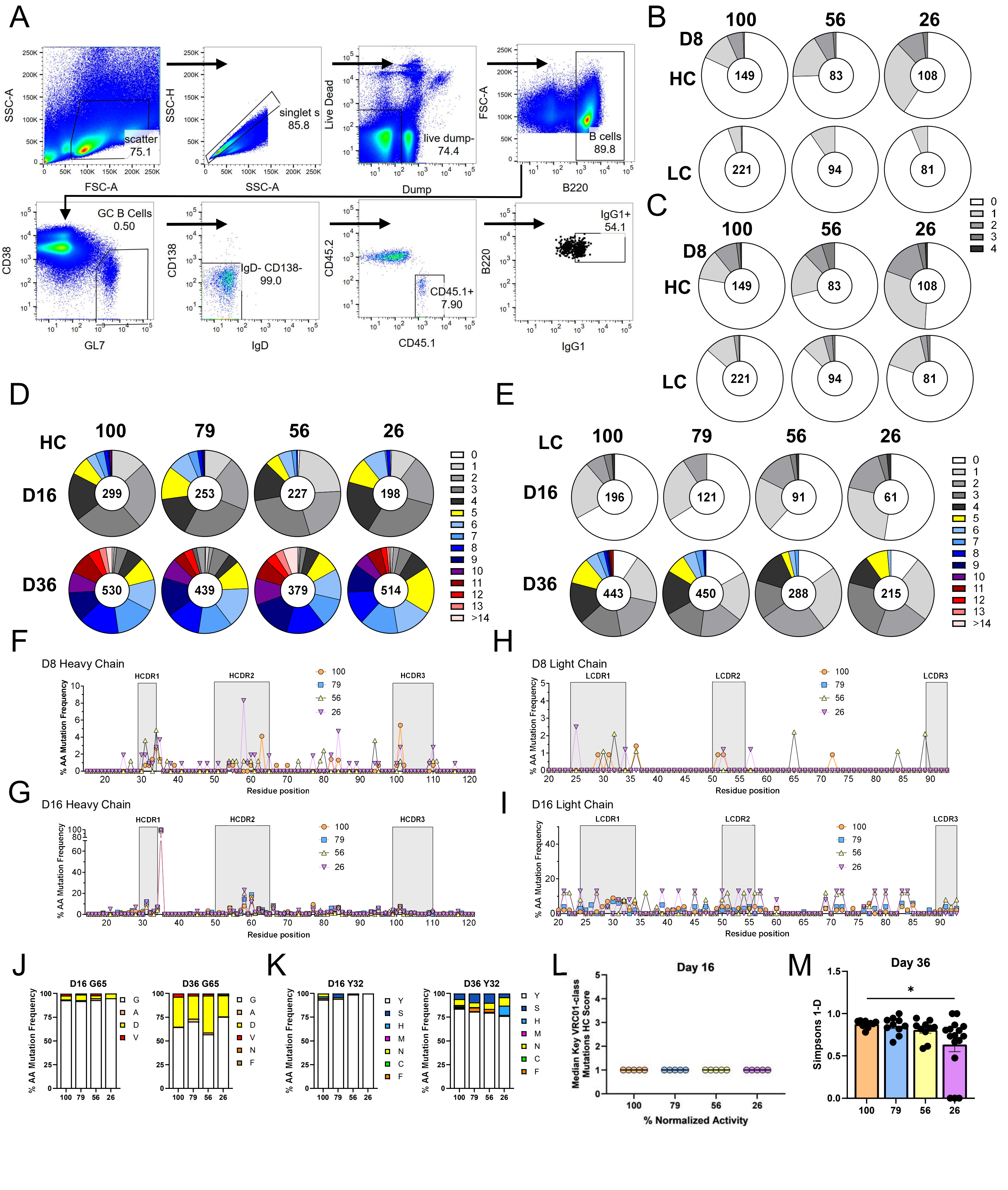

**Figure S2. Extended VRC01$^{gHL}$ mutation data, related to Figure 2 and Table S2. A)** Representative flow cytometry sort layout of VRC01$^{gHL}$ GC B cells from day 36. **B-C)** Circle charts represent the fraction of heavy chain (HC) and light chain (LC) sequences that acquired the indicated number of **(B)** amino acid mutations and **(C)** nucleotide mutations at day 8 post immunization. **D-E)** Circle charts represent the fraction of **(D)** HC and **(E)** LC sequences that acquired the indicated number of nucleotide mutations at days 16 and 36 post immunization. **B-E)** Total individual sequences are displayed in circle centers. **F-G)** Frequency of mutation at each residue in the VRC01$^{gHL}$ HC sequence at days **(F)** 8 and **(G)** 16. **H-I)** Frequency of mutation at each residue position in the VRC01$^{gHL}$ LC sequence at days **(H)** 8 and **(I)** 16. **J)** Frequency of G65 HC mutations at days 16 and 36. **K)** Frequency of Y32 LC mutation at days 16 and 36. **L)** Median values for key VRC01-class heavy chain residues from day 16 post prime. Dots represent individual mice with bars indicating the median of median values for each group. **M)** Simpson's index of diversity values for VRC01$^{gHL}$ GC BCR sequences on day 36. Dots indicate individual mice. Mean ± SEM. Kruskal-Wallis with Dunn's correction. *$p<0.05$.

### Supplementary Figure 3

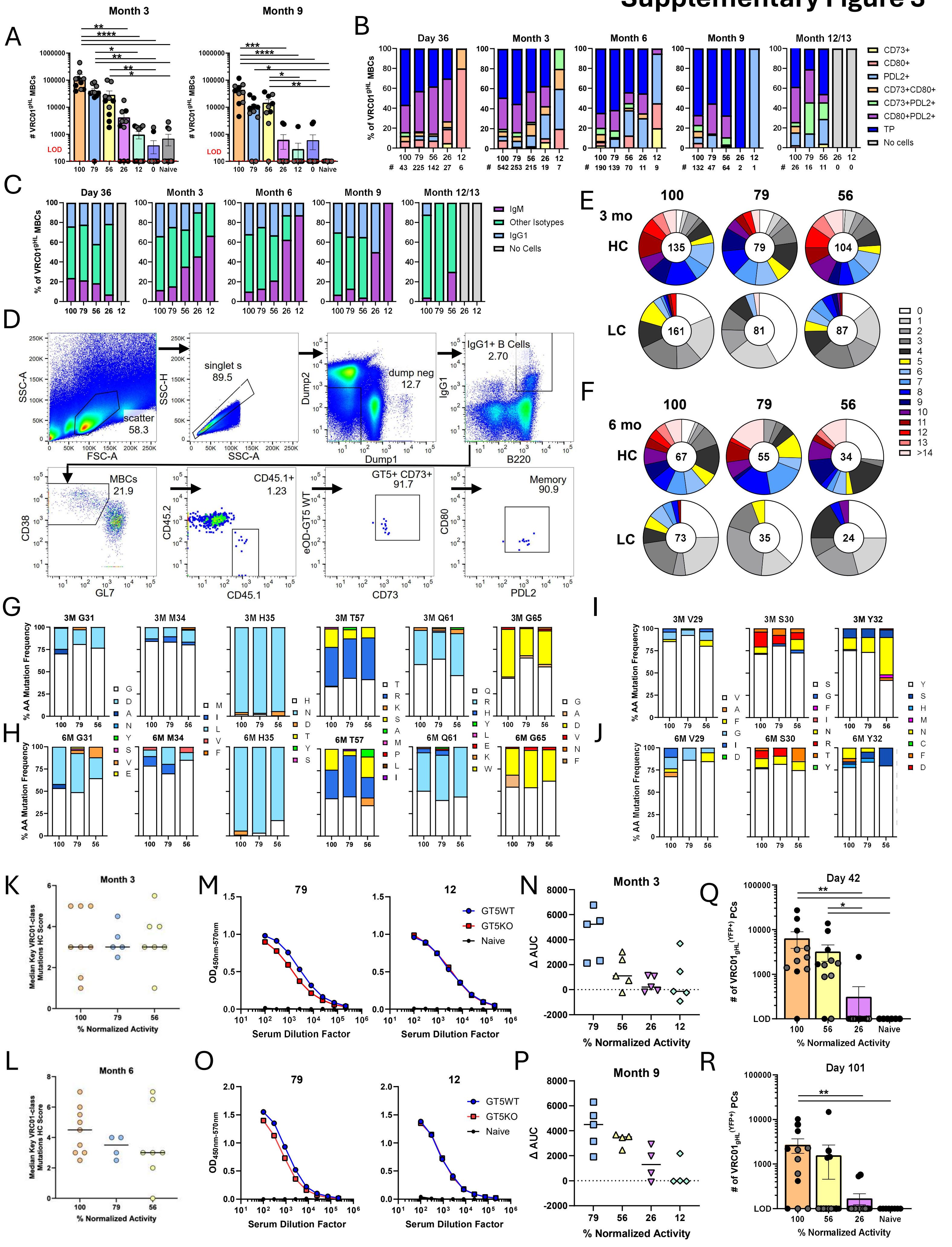

**Figure S3. Extended VRC01$^{gHL}$ MBC data, related to Figure 3. A)** Quantification of the absolute number of VRC01$^{gHL}$ MBCs. Isolated from whole spleen. **B)** Frequency of MBC phenotypes as determined by the three memory markers CD73, CD80 and PDL2 among total VRC01$^{gHL}$ MBCs. TP = triple positive for the three markers. **C)** Distribution of isotypes in VRC01$^{gHL}$ MBCs. **D)** Representative flow cytometry sort layout of VRC01$^{gHL}$ MBCs from month 6. **E-F)** Circle charts represent the fraction of heavy chain (HC) and light chain (LC) sequences from VRC01$^{gHL}$ MBCs that acquired the indicated number of nucleotide mutations at months 3 **(E)** and 6 **(F)** post immunization. Total individual sequences are displayed in circle centers. **G-H)** Frequency of key VRC01-class mutations in the heavy chain of VRC01$^{gHL}$ MBCs from months 3 **(G)** and 6 **(H)**. **I-J)** Frequency of key VRC01 mutations in the light chain of VRC01$^{gHL}$ MBCs from months 3 **(I)** and 6 **(J)**. **K-L)** Median values for key VRC01-class heavy chain residues from **(K)** 3 months and **(L)** 6 months post prime. Dots represent individual mice with bars indicating the median of median values for each group. **M)** Representative month 3 serum IgG ELISAs on mice immunized with 79% (left) and 12% (right) GT5 mosaic nanoparticles against GT5 WT (blue) and GT5 KO (red). **N)** CD4bs WT epitope specific responses from month 3 as determined by subtracting the KO area under the curve from the WT as shown in M. Shapes indicate individual mice. **O)** Representative month 9 serum IgG ELISAs on mice immunized with 79% (left) and 12% (right) GT5 mosaic nanoparticles against GT5 WT (blue) and GT5 KO (red). **P)** CD4bs WT epitope specific responses from month 9 as determined by subtracting the KO area under the curve from the WT as shown in O. Shapes indicate individual mice. **Q-R)** Quantification of the total number of VRC01$^{gHL}$ Blimp-1$^{YFP+}$ plasma cells at **(Q)** day 42 and **(R)** day 101 from bone marrow. **A, Q, R)** Mean ± SEM. Kruskal-Wallis with Dunn's correction. *$p<0.05$, **$p<0.01$, ***$p<0.001$, ****$p<0.0001$. LOD = limit of detection. Shades of grey dots indicate independent experiments.

### Supplementary Figure 4

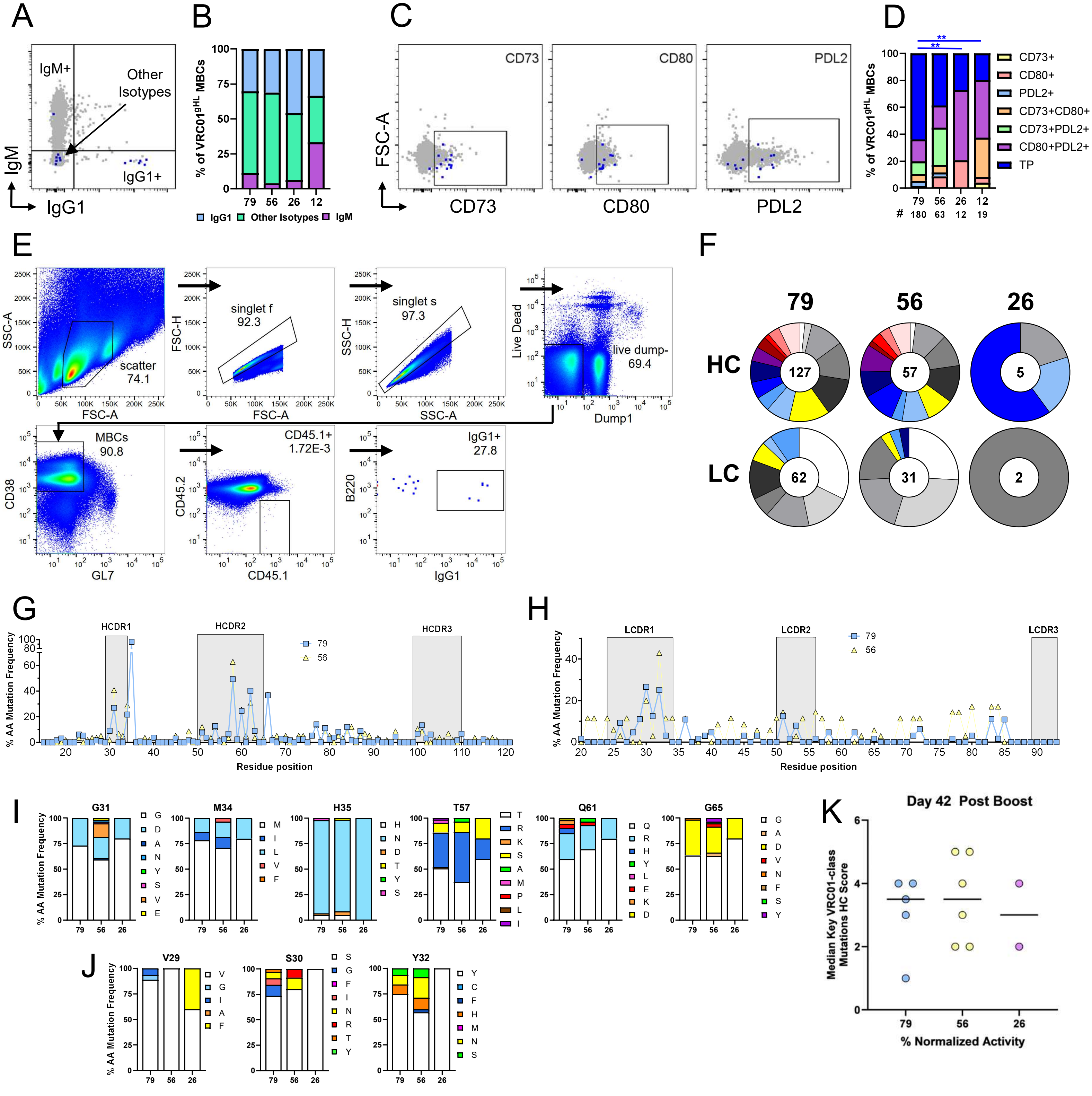

**Figure S4. Extended VRC01$^{gHL}$ MBC data from day 42 post-boost, related to Figure 4. A)** Representative flow cytometry plots of IgM+, IgG1+, and other isotypes of VRC01$^{gHL}$ MBCs (blue dots), overlaid on total endogenous MBCs from naïve control (grey). **B)** Isotype distribution of VRC01$^{gHL}$ MBCs from day 42 post boost. **C)** Representative flow cytometry plots of memory markers within VRC01$^{gHL}$ MBCs (blue dots), overlaid on total endogenous MBCs from naïve control (grey). **D)** Frequency of memory markers within VRC01$^{gHL}$ MBCs from day 42 post boost. The percent of marker triple positive (TP) MBCs were analyzed by Kruskal-Wallis with Dunn's correction. **$p<0.01$ **E)** Representative flow cytometry sort layout of VRC01$^{gHL}$ MBCs from day 42 post boost. **F)** Circle charts represent the fraction of heavy chain (HC) and light chain (LC) sequences from VRC01$^{gHL}$ MBCs that acquired the indicated number of nucleotide mutations at day 42 post boost. Total individual sequences are displayed in circle centers. **G-H)** Frequency of mutation at each residue position within the VRC01$^{gHL}$ **(G)** HC or **(H)** on day 42 post boost. **I)** Frequency of key VRC01 HC mutations in VRC01$^{gHL}$ MBCs. **J)** Frequency of key VRC01 LC mutations in VRC01$^{gHL}$ MBCs. **K)** Median values for key VRC01-class heavy chain residues from day 42 post boost. Dots represent individual mice with bars indicating the median of median values for each group.

### Supplementary Figure 5

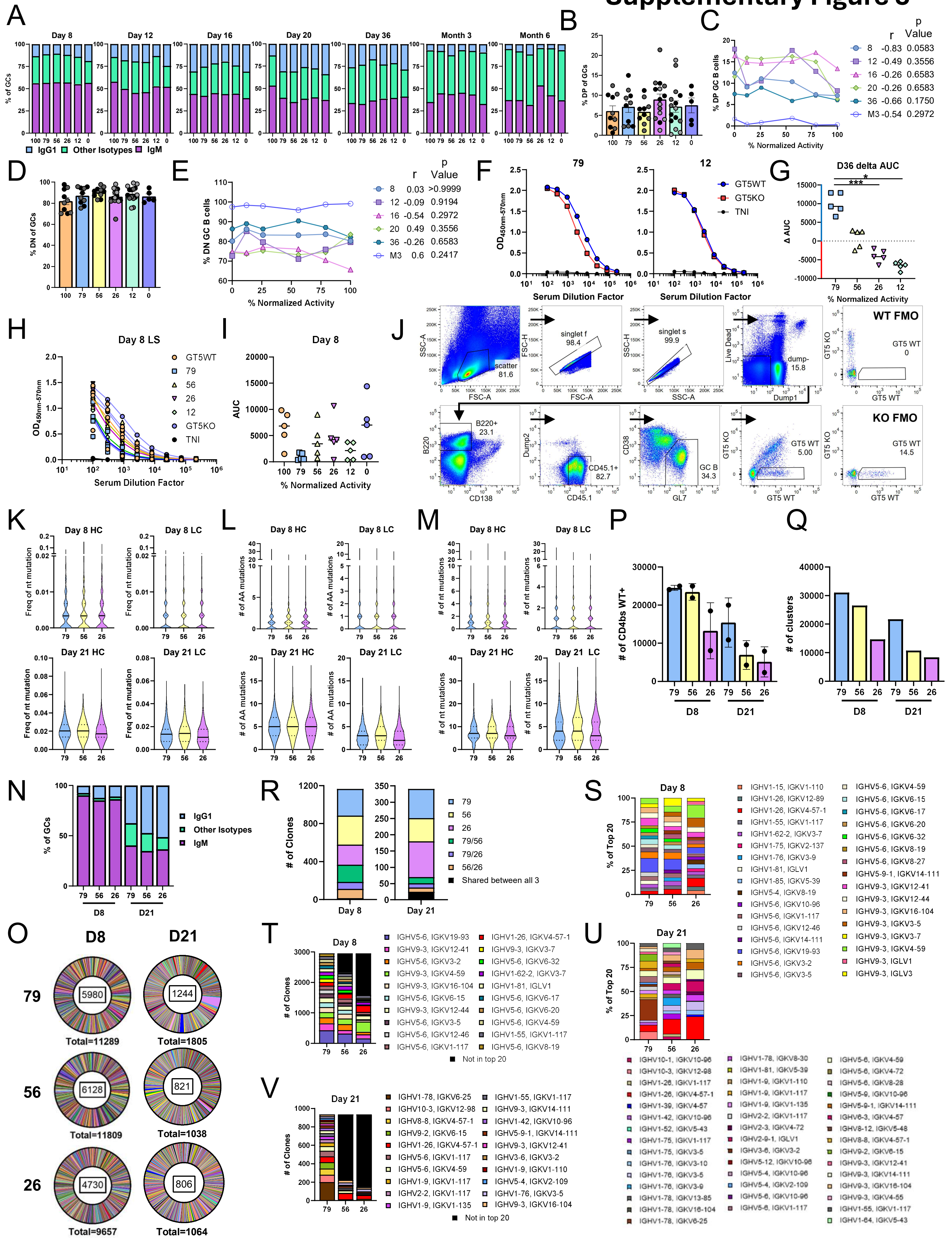

**Figure S5. Extended endogenous response data, related to Figure 5. A)** Isotype distribution of endogenous GC B cells from day 8 to month 6. **B)** Frequency of day 36 GC B cells double positive (DP) for binding both GT5 WT and KO probes. **C)** Line graph depicting average frequency of DP population in each group by timepoint. **D)** Frequency of day 36 GC B cells double negative (DN) for binding both GT5 WT and GT5 KO probes. **E)** Line graph depicting average frequency of DN population in each group by timepoint. **(C, E)** Spearman coefficient analysis for correlation between mosaic nanoparticle normalized activity and percent of GC B cells that were **(C)** DP or **(E)** DN. **F)** Representative day 36 serum IgG ELISAs on mice immunized with 79% (left) and 12% (right) GT5 mosaic nanoparticles against GT5 WT (blue) and GT5 KO (red). **G)** CD4bs WT epitope specific responses as determined by subtracting the KO area under the curve from the WT as shown in F. Shapes indicate individual mice. n= 5 mice/group. Kruskal-Wallis with Dunn's correction. *$p<0.05$, **$p<0.01$, ***$p<0.001$, ****$p<0.0001$. **H)** Day 8 serum IgG ELISA against lumazine synthase. **I)** Area under the curve of H. **J)** Representative flow cytometry sorting layout of WT CD4bs-specific endogenous (CD45.2+) GC B cells from days 8 and 21. WT CD4bs gating was determined by WT and KO FMO controls. **K)** Frequency of nucleotide mutations in heavy chains (HC) and light chains (LC) in endogenous GC B cells on days 8 and 21. **L)** Total number of aa mutations within HCs and LCs in endogenous GC B cells on days 8 and 21. **M)** Total number of nucleotide mutations in HCs and LCs in endogenous GC B cells on days 8 and 21. **N)** Isotype distribution of sorted cells. **O)** Circle graphs depicting the total number of unique clusters in recovered GC B cell clones at day 8 and day 21 post immunization. Center number indicates total number of clusters while total listed beneath indicates number of paired clones analyzed. **P)** Total number of WT CD4bs-specific cells collected on days 8 & 21. **Q)** Projected "absolute diversity" calculated by the number of unique clusters multiplied by total number of cells sorted per group. n= 8-9 mice/group. **R)** The distribution of clones that are unique or shared by GCs in response to mosaic nanoparticle immunization at days 8 and 21 as shown in Fig. 1K, M. **S)** Top 20 clonal families of each group at day 8. **T)** Quantification of top 20 clonal families present in mice immunized with the 79% mosaic nanoparticle and their representative frequency within the responding clones of mice immunized with 56% and 26% nanoparticles on day 8. Black bars indicate collated clonal families unique to 56% and 26% groups that were not seen in 79% group. **U)** Top 20 clonal families of each group at day 21. **V)** Quantification of top 20 clonal families present in mice immunized with the 79% mosaic nanoparticle and their respective frequency within the responding clones of mice immunized with 56% and 26% nanoparticles on day 21. Black bars indicate collated clonal families unique to 56% and 26% groups that were not seen in 79% group. **(B, D)** Mean ± SEM. Shades of grey dots indicate independent experiments.

### Supplementary Figure 6

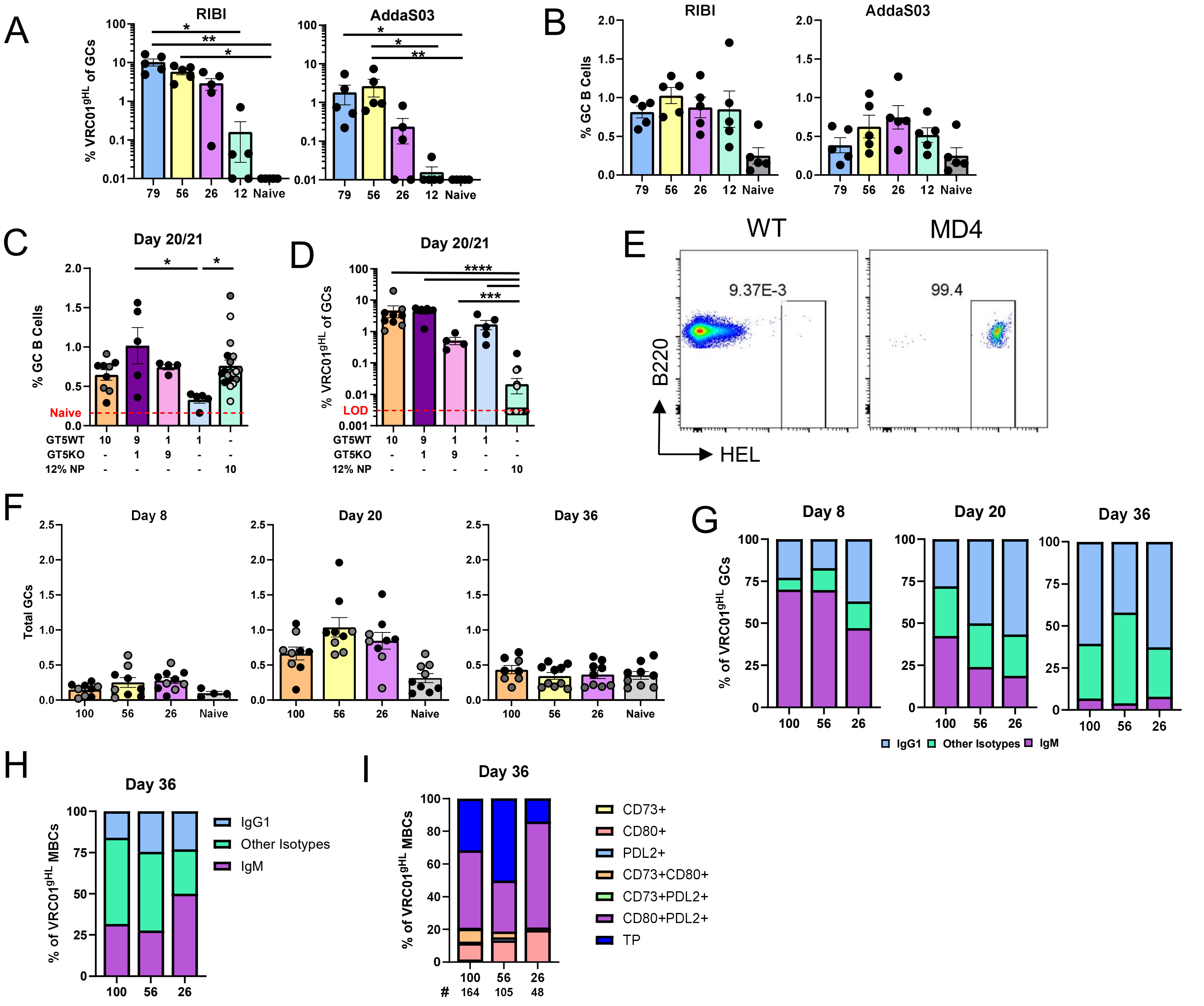

**Figure S6. Adjuvant, dosing, and extended MD4 model data, related to Figure 6. A)** Quantification of VRC01$^{gHL}$ frequency within GC B cells when immunogen is adjuvanted with (left) RIBI or (right) AddaS03. Kruskal-Wallis with Dunn's correction *p<0.05, **p<0.01. **B)** Frequency of total GC B cells when immunogen is adjuvanted with (left) RIBI or (right) AddaS03. **C)** Quantification of the frequency of total GC B cells on days 20/21 in mice immunized with one of the following: 10μg GT5-WT (day 20 & 21), 9μg/1μg GT5-WT/GT5-KO (day 21), 1μg/9μg GT5-WT/GT5-KO (day 21), 1μg GT5-WT (day 21), or 10μg of 12% (day 20) mosaic nanoparticle. 10μg GT5-WT and 10μg 12% mosaic nanoparticle data is re-graphed from Fig 1G. Kruskal-Wallis with Dunn's correction. *p<0.05. **D)** Quantification of the frequency of VRC01$^{gHL}$ within GCs on days 20/21 in mice that received the same immunizations as panel C. Mann Whitney U non-parametric tests were conducted, comparing each group to the 12% mosaic nanoparticle dose group. ***p<0.001, ****p<0.0001. **E)** Representative flow cytometry demonstrating hen-egg-lysozyme (HEL) specific binding of total B cells between wild-type and MD4 mice. **F)** Quantification of total GC frequency within MD4 mice immunized with mosaic nanoparticles. **G-H)** Isotype distribution of responding **(G)** GC and **(H)** memory B cells. **I)** Frequency of memory marker phenotypes in VRC01$^{gHL}$ MBCs. **(A-D, F)** Mean ± SEM. Shades of grey dots indicate independent experiments.

### Supplementary Figure 7

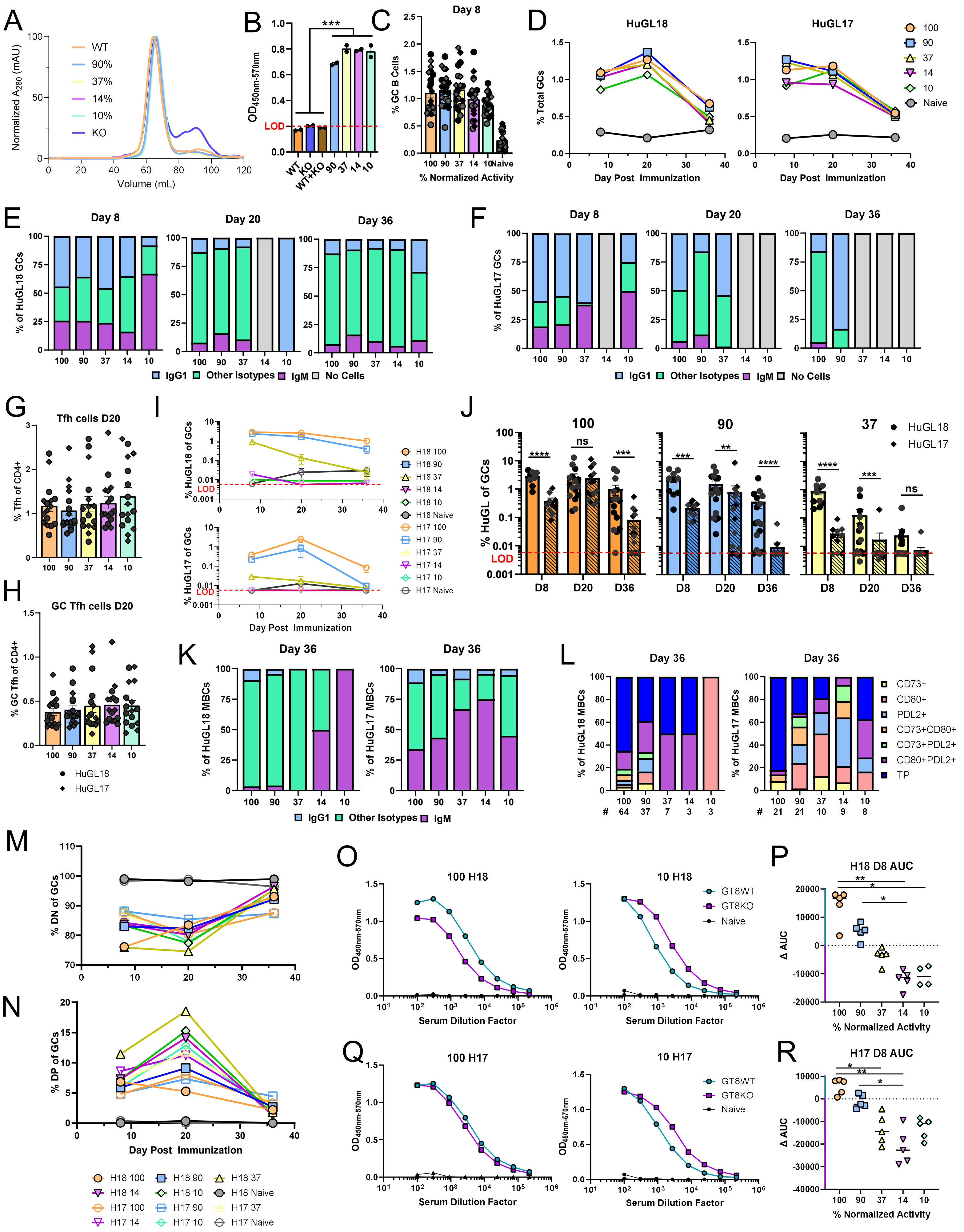

**Figure S7. Characterization of eOD-GT8 mosaic nanoparticles and extended HuGL18 and HuGL17 data, related to Figure 7 and Table S4. A)** Normalized SEC profiles of eOD-GT8 mosaic nanoparticles. **B)** Results of ELISA using schematic in Fig. S1A, demonstrating that only GT8 mosaic nanoparticles bind both CD4bs WT and KO specific antibodies, compared to fully WT or KO 60mer nanoparticles. Each dot indicated technical replicate. Mann-Whitney U test comparing pooled non-mosaic particle to mosaic particle groups. ***p<0.001. **C)** Frequency of total GC B cells in response to HuGL transfer and GT8 nanoparticle immunization. **D)** Frequency of total GC B cells over time in (left) HuGL18 and (right) HuGL17 recipients as shown in C. **E)** Isotype distribution in HuGL18 GC B cells. **F)** Isotype distribution in HuGL17 GC B cells. **G)** Frequency of Tfh (PD1+/CXCR5+/Foxp3-) among CD4+ T cells (SSL/B220-/CD4+) in spleens of immunized mice. **H)** Frequency of GC Tfh (CD90.2- Tfh) among CD4+ T cells in spleens of immunized mice. **I)** Frequency of HuGL18 (top) or HuGL17 (bottom) GC B cells over time. **J)** Direct comparison of HuGL18 and HuGL17 frequencies in GCs over time in mice receiving either 100 (left), 90 (middle) or 37 (right) mosaic particles. Mann Whitney U non-parametric tests were conducted comparing HuGL18 to HuGL17 at each timepoint for each group. ns = not significant, **p<0.01, ***p<0.001, ****p<0.0001. **(I-J)** Data re-graphed from Fig. 7D, G. LOD = limit of detection. **K)** Frequency of memory marker phenotypes on (left) HuGL18 and (right) HuGL17. **L)** Isotype distribution of (left) HuGL18 and (right) HuGL17 MBCs. **M)** Frequency of GC B cells double negative (DN) for binding GT8 WT and KO probes on days 8-36. **N)** Frequency of GC B cells double positive (DP) for binding GT8 WT and KO probes on days 8-36. **O)** Representative day 8 serum IgG ELISAs on HuGL 18 mice immunized with 100% (left) and 10% (right) GT8 mosaic nanoparticles against GT8 WT (teal) and GT8 KO (purple). **P)** CD4bs WT epitope specific responses from HuGL18 recipients on day 8 as determined by subtracting the KO area under the curve from the WT as shown in O. Shapes indicate individual mice. **Q)** Representative day 8 serum IgG ELISAs on HuGL 17 mice immunized with 100% (left) and 10% (right) GT8 mosaic nanoparticles against GT8 WT (teal) and GT8 KO (purple). **R)** CD4bs WT epitope specific responses from HuGL17 recipients on day 8 as determined by subtracting the KO area under the curve from the WT as shown in Q. Shapes indicate individual mice. **P, R)** Kruskal-Wallis with Dunn's correction. *p<0.05, **p<0.01. **(C, G-H, J)** Circles indicate HuGL18 and diamonds indicate HuGL17. **(B-C, G-H, J)** Mean ± SEM. Shades of grey indicate independent experiments.

**Table S1. Co-transfection ratios and KinExa data for eOD-GT5 mosaic nanoparticles.**

| Name | %WT plasmid in cotransfection | $K_D$ for VRC01 (M) | KinExa Titrant % Activity | Normalized % Activity |
|---|---|---|---|---|
| eOD-GT5 60mer$^{100}$ prep1 | 100% | 2.06E-10 | 43 | 100 |
| eOD-GT5 60mer$^{79}$ prep1 | 90% | 1.91 E-10 | 34 | 79 |
| eOD-GT5 60mer$^{56}$ prep1 | 50% | 1.82 E-10 | 24 | 56 |
| eOD-GT5 60mer$^{26}$ prep1 | 20% | 9.70 E-11 | 11 | 26 |
| eOD-GT5 60mer$^{12}$ prep1 | 10% | 1.29 E-10 | 5 | 12 |
| eOD-GT5 60mer$^{0}$ prep1 | 0% | n.b. | 0 | 0 |
| eOD-GT5 60mer$^{100}$ prep2 | 100% | 1.84 E-10 | 38 | 100 |
| eOD-GT5 60mer$^{79}$ prep2 | 90% | 1.21 E-10 | 31 | 82 |
| eOD-GT5 60mer$^{56}$ prep2 | 50% | 1.95 E-10 | 20 | 53 |
| eOD-GT5 60mer$^{26}$ prep2 | 20% | 1.63 E-10 | 9 | 24 |
| eOD-GT5 60mer$^{12}$ prep2 | 10% | 1.59 E-10 | 5 | 13 |
| eOD-GT5 60mer$^{0}$ prep2 | 0% | n.b. | 0 | 0 |

**Table S2. Number of amino acid mutations in VRC01$^{gHL}$ GC and MBC sequences.**

| Timepoint | NP | Median | Mean | *E* | *n* sequences | *n* mice |
|---|---|---|---|---|---|---|
| **Heavy Chain** | | | | | | |
| D8[a] | 100 | 0 | 0.26 | 1 | 149 | 4 |
| | 56 | 0 | 0.35 | 1 | 83 | 2 |
| | 26 | 0 | 0.55 | 2 | 108 | 8 |
| D16[a] | 100 | 2 | 2.41 | 1 | 299 | 5 |
| | 79 | 2 | 2.59 | 1 | 253 | 5 |
| | 56 | 2 | 2.14 | 1 | 227 | 5 |
| | 26 | 2 | 2.45 | 1 | 221 | 5 |
| D36[a] | 100 | 6 | 5.65 | 2 | 525 | 10 |
| | 79 | 5 | 5.11 | 2 | 439 | 10 |
| | 56 | 6 | 5.94 | 2 | 371 | 10 |
| | 26 | 5 | 4.85 | 5 | 502 | 22 |
| 3 mo[b] | 100 | 6 | 6.02 | 2 | 135 | 8 |
| | 79 | 4 | 4.97 | 2 | 79 | 7 |
| | 56 | 5.5 | 5.81 | 2 | 104 | 9 |
| 6 mo[b] | 100 | 6 | 6.06 | 2 | 67 | 9 |
| | 79 | 5 | 5.77 | 2 | 53 | 6 |
| | 56 | 6 | 5.97 | 2 | 36 | 8 |
| D42PB[b] | 79 | 4 | 4.33 | 2 | 127 | 10 |
| | 56 | 4 | 5.04 | 2 | 57 | 10 |
| | 26 | 6 | 5.6 | 1 | 5 | 2 |
| **Light Chain** | | | | | | |
| D8[a] | 100 | 0 | 0.6 | 1 | 221 | 4 |
| | 56 | 0 | 0.1 | 1 | 94 | 2 |
| | 26 | 0 | 0.06 | 2 | 81 | 8 |
| D16[a] | 100 | 0 | 0.55 | 1 | 301 | 5 |
| | 79 | 0 | 0.32 | 1 | 139 | 5 |
| | 56 | 0 | 0.45 | 1 | 91 | 5 |
| | 26 | 0 | 0.44 | 1 | 61 | 5 |
| D36[a] | 100 | 2 | 2.12 | 2 | 443 | 10 |
| | 79 | 2 | 1.98 | 2 | 450 | 10 |
| | 56 | 1 | 1.6 | 2 | 288 | 10 |
| | 26 | 2 | 1.61 | 4 | 215 | 18 |
| 3 mo[b] | 100 | 2 | 2.25 | 2 | 161 | 9 |
| | 79 | 0 | 1.19 | 1 | 81 | 5 |
| | 56 | 2 | 2.21 | 2 | 87 | 8 |
| 6 mo[b] | 100 | 1 | 1.82 | 2 | 73 | 10 |
| | 79 | 1 | 1.17 | 2 | 35 | 5 |
| | 56 | 1.5 | 1.5 | 2 | 24 | 8 |
| D42PB[b] | 79 | 1 | 1.58 | 2 | 62 | 9 |
| | 56 | 1 | 1.55 | 2 | 31 | 8 |
| | 26 | 2 | 1.33 | 1 | 3 | 2 |

**NP** = Nanoparticle normalized percent binding activity
**Median** = Median number of amino acid mutations
**Mean** = Average number of amino acid mutations
***E*** = Number of experiments
***n* sequences** = Number of sequences analyzed
***n* mice** = Number of mice
[a] Germinal Center B cells
[b] Memory B cells

#### Table S3. Simpon's Index of Diversity for VRC01$^{gHL}$ GC and MBC BCR sequencing.

| Timepoint | Group | M1 | M2 | M3 | M4 | M5 | M6 | M7 | M8 | M9 | M10 | M11 | M12 | M13 | M14 | M15 | M16 | Mean | SD |
|---|---|---|---|---|---|---|---|---|---|---|---|---|---|---|---|---|---|---|---|
| Day 16 | 100 | 0.525 | 0.645 | 0.330 | 0.394 | 0.798 | - | - | - | - | - | - | - | - | - | - | - | 0.538 | 0.189 |
| | 79 | 0.154 | 0.614 | 0.621 | 0.554 | 0.417 | - | - | - | - | - | - | - | - | - | - | - | 0.472 | 0.196 |
| | 56 | 0.472 | 0.493 | 0.000 | 0.000 | 0.242 | - | - | - | - | - | - | - | - | - | - | - | 0.241 | 0.241 |
| | 26 | 0.564 | 0.400 | 0.000 | 0.000 | 0.200 | - | - | - | - | - | - | - | - | - | - | - | 0.233 | 0.248 |
| Day 36 | 100 | 0.835 | 0.858 | 0.886 | 0.912 | 0.879 | 0.885 | 0.901 | 0.910 | 0.917 | 0.781 | - | - | - | - | - | - | 0.876 | 0.042 |
| | 79 | 0.917 | 1.000 | 0.733 | 0.819 | 0.871 | 0.662 | 0.947 | 0.834 | 0.922 | 0.855 | - | - | - | - | - | - | 0.856 | 0.101 |
| | 56 | 0.588 | 0.832 | 0.818 | 0.616 | 0.857 | 0.786 | 0.868 | 0.921 | 0.945 | 0.838 | - | - | - | - | - | - | 0.807 | 0.118 |
| | 26 | 0.817 | 0.476 | 0.848 | 1.000 | 0.742 | 0.000 | 0.733 | 0.806 | 0.867 | 0.733 | 0.800 | 0.667 | 0.000 | 1.000 | 0.679 | 0.000 | 0.636 | 0.339 |
| 3 mo | 100 | 1.000 | 0.000 | 1.000 | 0.778 | 0.500 | 0.906 | 0.919 | 1.000 | - | - | - | - | - | - | - | - | 0.763 | 0.352 |
| | 79 | 1.000 | 0.889 | 0.800 | 0.909 | 0.861 | - | - | - | - | - | - | - | - | - | - | - | 0.892 | 0.073 |
| | 56 | 0.000 | 1.000 | 0.733 | 0.933 | 0.000 | 0.918 | 0.667 | - | - | - | - | - | - | - | - | - | 0.607 | 0.431 |
| 6 mo | 100 | 0.000 | 0.000 | 0.000 | 0.833 | 0.000 | 0.929 | 0.700 | 0.800 | 0.000 | - | - | - | - | - | - | - | 0.362 | 0.434 |
| | 79 | 0.000 | 0.000 | 1.000 | 0.857 | - | - | - | - | - | - | - | - | - | - | - | - | 0.464 | 0.539 |
| | 56 | 0.000 | 0.000 | 1.000 | 0.833 | 0.000 | 0.833 | 0.000 | - | - | - | - | - | - | - | - | - | 0.381 | 0.478 |
| D42 PB | 79 | 0.781 | 0.000 | 0.000 | 0.643 | 0.833 | - | - | - | - | - | - | - | - | - | - | - | 0.451 | 0.418 |
| | 56 | 0.733 | 1.000 | 0.000 | 0.000 | 0.000 | 0.000 | - | - | - | - | - | - | - | - | - | - | 0.289 | 0.455 |
| | 26 | 0.000 | 0.000 | - | - | - | - | - | - | - | - | - | - | - | - | - | - | 0.000 | 0.000 |

**Table S4. Simpon's Index of Diversity for Endogenous CD4bs WT GC B cells**

| | 79 | 56 | 26 |
|---|---|---|---|
| **Day 8** | 0.950293 | 0.934886 | 0.915961 |
| **Day 21** | 0.994193 | 0.998252 | 0.998589 |

**Table S5. Quantification of affinity from antibodies derived from cultured GC B cells.**

| Timepoint | Day 8 | | | Day 21 | | |
|---|---|---|---|---|---|---|
| % Normalized Activity | 79 | 56 | 26 | 79 | 56 | 26 |
| *N* Mice | 4 | 4 | 3 | 4 | 5 | 4 |
| *N* mAbs tested (total) | 189 | 153 | 150 | 166 | 177 | 179 |
| WT $K_D$s <100 µM (non-binders) (Excluded) | 89 | 80 | 113 | 53 | 79 | 80 |
| KO $K_D$s <100 µM (cross reactive) (Excluded) | 45 | 34 | 14 | 66 | 50 | 64 |
| *N* WT-only binders | **55** | **39** | **23** | **47** | **48** | **35** |
| WT-only binders median $K_D$ (µM) | 12 | 7.75 | 7.21 | 1.026 | 4.09 | 11.3 |
| WT-only binders geometric mean $K_D$ (µM) | 8.02 | 5.8 | 6.33 | 0.85 | 1.64 | 6.98 |
| Geometric mean fold change from day 8 | | | | **9.44** | **3.51** | **0.91** |

**Table S6. Co-transfection ratios and KinExa data for eOD-GT8 mosaic nanoparticles.**

| Name | %WT plasmid in cotransfection | $K_D$ for GL-VRC01 (M) | KinExa Titrant % Activity | Normalized % Activity |
|---|---|---|---|---|
| eOD-GT8 60mer$^{100}$ | 100% | 2.40E-11 | 68 | 100 |
| eOD-GT8 60mer$^{90}$ | 90% | 2.70E-11 | 61 | 90 |
| eOD-GT8 60mer$^{37}$ | 50% | 2.00E-11 | 25 | 37 |
| eOD-GT8 60mer$^{14}$ | 20% | 1.40E-11 | 10 | 14 |
| eOD-GT8 60mer$^{10}$ | 10% | 1.50E-11 | 7 | 10 |
| eOD-GT8 60mer$^{0}$ | 0% | n.b. | 0 | 0 |
